# Nuclear dualism without extensive DNA elimination in the ciliate *Loxodes magnus*

**DOI:** 10.1101/2023.11.09.566212

**Authors:** Brandon K. B. Seah, Aditi Singh, David E. Vetter, Christiane Emmerich, Moritz Peters, Volker Soltys, Bruno Huettel, Estienne Swart

## Abstract

Ciliates are unicellular eukaryotes with two distinct kinds of nuclei in each cell: transcriptionally active somatic macronuclei (MAC) and silent germline micronuclei (MIC). In the best-studied model species, both nuclei can divide asexually, but only germline MICs participate in meiosis, karyogamy, and development into new MACs. During MIC-to-MAC development, thousands of mobile element relics in the germline, called internally eliminated sequences (IESs), are excised. This genome editing enables IESs to persist by shielding them from somatic natural selection. Editing itself is a costly, time-consuming process, hypothetically maintained by evolutionary addiction. *Loxodes magnus* and its relatives (class Karyorelictea) are cytologically unusual because their MACs do not divide asexually, but must develop anew from mitotically generated MIC copies every cell division. Here, we report that *Loxodes* genome development is also unconventional. We found no canonical germline-limited IESs in *Loxodes* despite careful purification and long-read sequencing of MICs and MACs. The k-mer content of these nuclei overlapped, and indels found by read mapping were consistent with allele variants rather than IESs. Two other hallmarks of genome editing—domesticated DDE-family transposases and editing-associated small RNAs—were also absent. Nonetheless, histone marks, nucleosome and DNA N6-methyladenosine distributions in vegetative *Loxodes* cells are consistent with actively transcribed MACs and inactive MICs, like other ciliates. Both genomes, not only the MIC, were large and replete with retrotransposon sequences. Given the costs associated with genome editing, we hypothesize that karyorelicteans like *Loxodes* have lost or streamlined editing during MIC-to-MAC development, and have found a way out of the addictive cycle.

## Introduction

Ciliate genomes undergo profound changes during development. Each cell has two types of nuclei (nuclear dualism): smaller, germline micronuclei (MICs) and larger, somatic macronuclei (MACs). Both divide asexually, but during sexual reproduction, only MICs undergo meiosis and karyogamy to form a diploid zygotic nucleus, which develops into new MACs that replace the old MACs.^1^ During this development, a significant fraction of the MIC genome is eliminated^2–4^, largely composed of repetitive elements like microsatellites, minisatellites, and transposons; known MIC genomes are hence ∼10 to 450 Mbp larger than MACs (∼40 to 100 Mbp length).^5–13^ The remaining DNA is amplified to 10s to 10000s of copies depending on species,^1,2,14,15^ to form the mature “ampliploid” MAC.^3^ MIC chromosomes are also fragmented into shorter MAC DNA molecules during development;^1^ the degree also varies between species, with an extreme of kilobase-sized single-gene “nanochromosomes” in spirotrichs.^13,16–18^

Most of the eliminated DNA comprises “internally eliminated sequences” (IESs), where flanking segments are joined after excision. Their length, placement, and content are variable,^1,14^ e.g. mostly <100 bp in *Paramecium* but ∼10 kbp in *Tetrahymena*.^6,7^ IESs are thought to originate from DNA transposons, and the excisases that remove them evolved from DDE-family DNA transposases.^7–9,19–22^ Excision is proposed to be guided by development-specific small RNAs of different classes.^23–27^ Genome editing removes mobile elements from the MAC, as a result, they are not exposed to natural selection and tend to accumulate in MIC genomes over time.^8,22,28^

Both the chemical nature of DNA and chromatin also differ between the two nuclei: histone variants may be MIC- or MAC-specific;^29–31^ nucleosomes are distinctly phased relative to gene features in MACs but not MICs;^32^ >1% of adenosines in MAC DNA have N6-methyl-deoxyadenosine (6mA) modification vs. negligible levels in MICs.^33–36^

A possible exception to aspects of this general picture is the class Karyorelictea (Figure 1A, 1B), whose development differs from that of model ciliates in two major ways: (i) their MACs cannot divide and always develop from MIC precursors, even during asexual division,^37^ and (ii) karyorelict MACs are less amplified than other ciliates (“paradiploid” vs. ampliploid), e.g. in the karyorelict *Loxodes magnus*, the DNA content in MACs is only about twice that of MICs.^38^ Karyorelicts were formerly considered a “basal” group with a “primitive” form of nuclear development.^37,39^ However, molecular phylogenies now show that karyorelicts are not the earliest-diverging branch but in fact sister to the Heterotrichea,^40–42^ which have dividing, ampliploid MACs and at least one genus with extensive genome editing like other ciliates,^9,12^ so non-dividing paradiploid MACs must be a derived character of karyorelicts.

**Figure 1.**
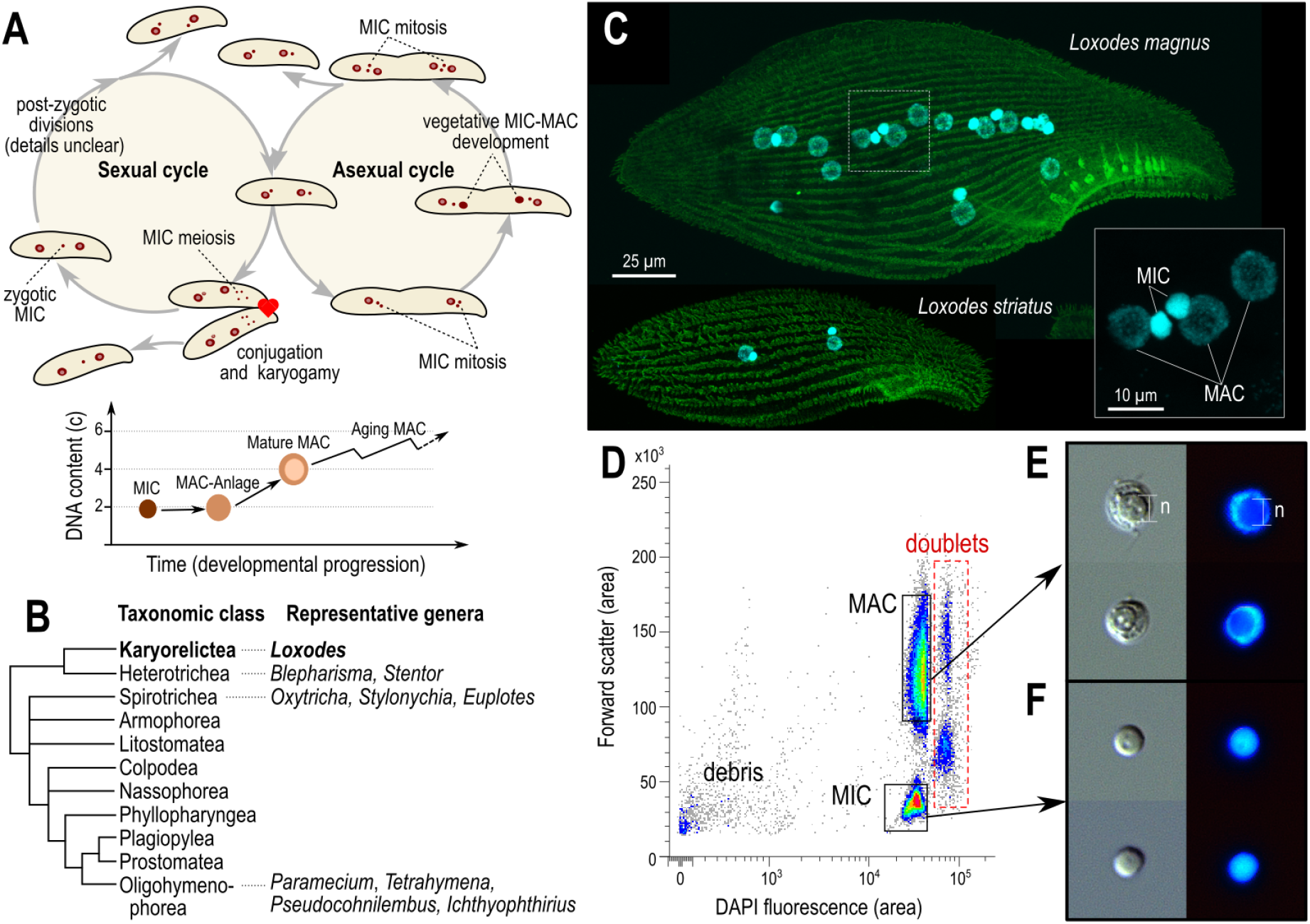
*Loxodes* nuclei purification (A) Schematic of nuclei in *Loxodes striatus* sexual and asexual cycles (above), adapted from ^37,71,72^; diagram of DNA content in Loxodes nuclei at different developmental stages (below), adapted from ^38^. **(B)** Diagrammatic tree of ciliate classes (after ^66^, branch lengths arbitrary) with genera used as laboratory models. **(C)** Confocal scanning fluorescence micrographs of *Loxodes magnus* and *L. striatus* cells (maximum-intensity projections): green, alpha-tubulin secondary immunofluorescence; cyan, DAPI staining of nuclei; inset, detail of nuclei from *L. magnus*. **(D)** Representative flow cytometric scatter plot of forward scatter vs. DAPI fluorescence for *L. magnus* cell lysate, with gates for MAC and MIC defined for flow sorting. **(E)** and **(F)** MAC and MIC respectively after sorting, imaged with differential interference contrast (left) and DAPI stain (right); each subpanel width 10 µm. The nucleolus (“n”) is a spherical region less densely stained with DAPI in panel C.

What consequences does this decoupling of MIC-to-MAC development from the sexual cycle have for genome development and evolution? Given that karyorelicts must undergo this development with every cell division, we hypothesized that if IESs are present they would also need to be excised every division. We therefore sequenced both genomes from the karyorelict *Loxodes* (Figure 1C) to compare their sequence content and architecture.

Unexpectedly we did not detect classical IESs, suggesting that their genomes are on a different evolutionary trajectory from all other ciliates studied to date.

## Results

### Physical purification of *Loxodes* MICs and MACs

Two distinct clusters corresponding to MACs and MICs were observed in fluorescence activated nuclear sorting of DAPI-stained cell lysates of *Loxodes magnus* (Figures 1D, 1E, 1F, S1A) and *L. striatus* (Figure S1B). Sorted nuclear purity was also verified by distinct histone marks in the two sorted nuclei populations, and presence vs. absence of the 6mA base modification (Results: “*Loxodes* MACs have characteristics of active chromatin”).

Sorted samples were largely free of bacterial contamination: in short read libraries prepared from sorted MIC and MAC libraries, at most 1.3% of reads mapping to SSU rRNA sequences were classified as bacterial or archaeal (Figure S2).

### *Loxodes* MIC and MAC genomes have similar k-mer composition

We first compared the composition of short subsequences of defined length, known as k-mers, in the unassembled short read *Loxodes* MIC and MAC genome libraries (k=21 nt). Most k-mers observed were shared by both libraries. Of k-mers with combined frequency ≥5× in *L. magnus*, only 3.3% were unique to one or the other library, whereas 93% were observed ≥5× in each library. Unique k-mers did not show discernible frequency peaks (Figure 2A), nor was there an obvious cluster of k-mers with different coverage between the two libraries (Figure 2B), contrary to what would be expected if much of the genome was MIC-limited like in other ciliates (Supplementary Results 1), or if there was differential amplification of specific loci, as previously proposed.^38,43^ There was no evidence for amplification of the rRNA locus in particular (Supplementary Results 2).

**Figure 2.**
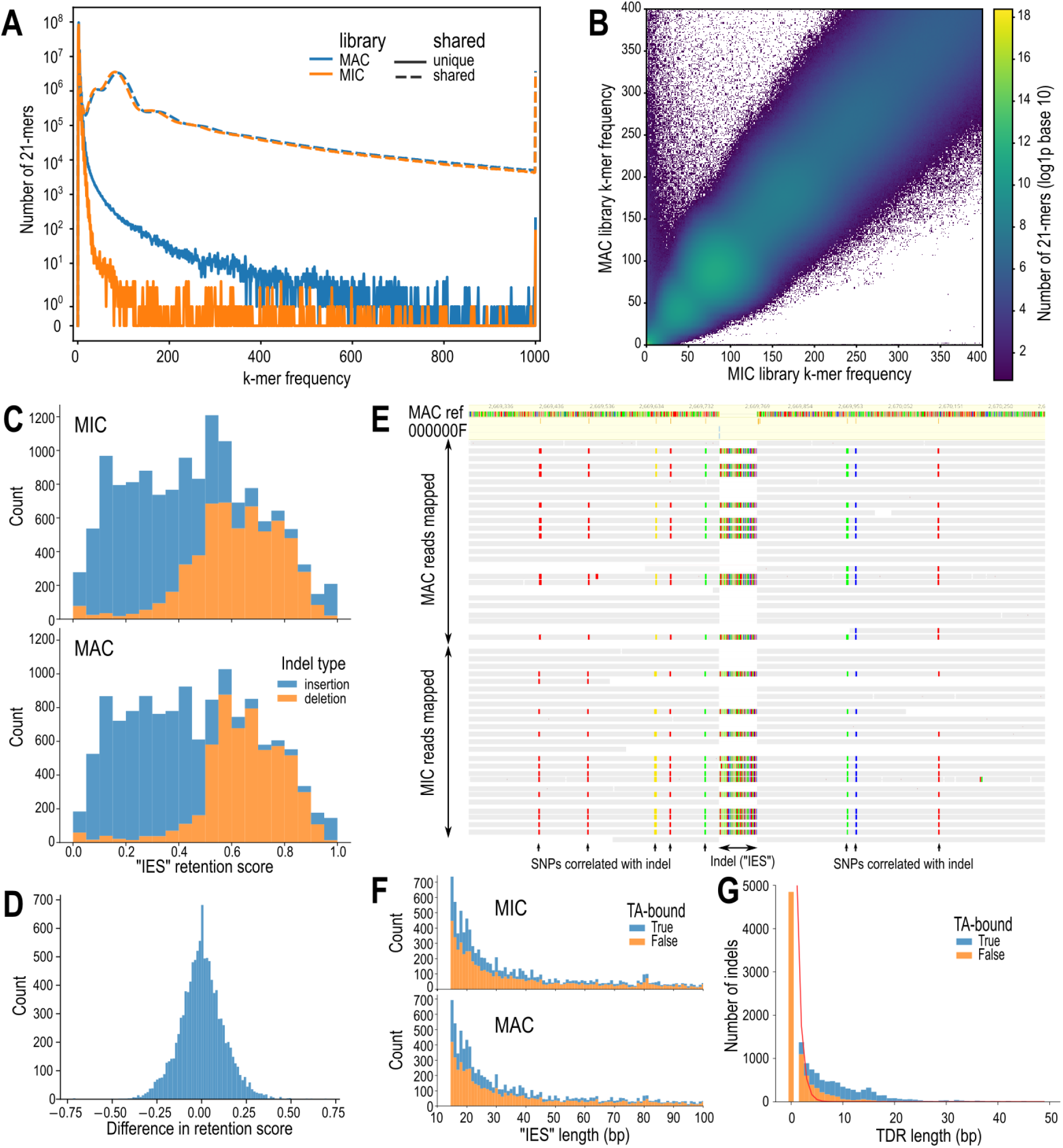
Screening for IESs in *Loxodes magnus*. (A) k-mer multiplicity plot for shared (dashed lines) vs. unique (solid lines) 21-mers in MAC (blue) and MIC (orange) sequence libraries of *L. magnus*. **(B)** Heatmap comparing frequency of genomic 21-mers in MIC vs. MAC of *L. magnus*; color scale represents log (1 + number of k-mers); axes truncated at 400x frequency. **(C)** Histograms of relative coverage (“retention score”) for putative “IESs” (indels) predicted by an IES detection pipeline from MIC and MAC sorted nuclei long-read libraries. **(D)** Histogram of differences in retention scores between MIC and MAC libraries for putative “IESs”. **(E)** Example of PacBio HiFi long reads (horizontal bars) from MIC and MAC libraries mapped to MAC reference genome (colored bar, top), containing an “IES” indel correlated with SNPs; colored positions in reads represent bases different from reference. **(F)** Length histograms of indel polymorphisms; colored by whether they are bound by TA-containing tandem repeats; x-axis truncated at 100 bp. **(G)** Lengths of tandem direct repeats bounding indel polymorphisms (bars), compared to expected lengths assuming random sequence (red line).

k-mer frequency spectra of both nuclei each showed a main coverage peak (∼85×), a heterozygosity peak (∼40×), and additional peaks (∼170× and ∼430×) which suggested some degree of genome duplication or paralogy. The spectra were long-tailed; 0.68% of k-mers had ≥1000× frequency, representing high-copy-number repeat elements in the genome. Low frequency k-mers were likely sequencing errors (18% singletons, 56% with combined frequency <5×) (Figure 2A). Similar genome sizes (262 Mbp MIC, 261 Mbp MAC) and heterozygosity (0.60% and 0.59%) were predicted from model-fitting of k-mer coverage spectra peaks, although these do not account for high copy repeats. *L. striatus* k-mer spectra showed similar patterns (Figure S3, S4).

### Classical IESs not detected in Loxodes magnus MIC genome

We next attempted to detect IESs in *L. magnus* by calling indels from error-corrected long reads (PacBio HiFi) mapped to the MAC reference assembly. If there are MIC-specific IESs, they should present in reads as insertions relative to the MAC reference, and the fraction of reads bearing the insert (“retention score”) should be significantly higher in the MIC than the MAC (Supplementary Results 3). Slightly more candidate “IESs” were called from the MIC (13,734) vs. MAC (12,897), of which 10,992 were predicted in both. However, the mean retention scores per library were similar (0.45 for MIC, 0.46 for MAC) (Figure 2C), and retention scores of shared “IESs” were not significantly different between the two libraries (Figure 2D, Wilcoxon signed-rank test, one-sided for higher score in MIC, *p*=0.29). The relative proportion of insertions vs. deletions was similar between MIC and MAC libraries (Figure 2C), contrary to the expectation of more inserts in the MIC library. Indels that were both unique to the MIC library and with high retention score (>0.9), as would be expected of true IESs, were few in number (40) and located in regions of low coverage (mean 4.2×), and were hence probably mispredictions due to insufficient coverage.

“IESs” from *L. magnus* were instead consistent with allelic indel polymorphisms, because inserts had a coverage of about 50% and the presence of an insert in a read was usually correlated with single nucleotide polymorphisms, regardless of the nucleus type (Figure 2E, Figure S5, Supplementary Results 3). Their length distribution also lacked specific peaks, like in other ciliates, but simply sloped downwards from the lower length cutoff (Figure 2F). Nonetheless, more indels were bound by terminal direct repeats (TDR) than expected by chance, especially TDRs that contain TA-sequence submotifs (Figure 2G), and hence could have originated from mobile elements.

### Both *Loxodes magnus* nuclear genomes are rich in tandem and interspersed repeats

*Loxodes magnus* genome assemblies from long reads were large (MIC 848 Mbp, MAC 706 Mbp), but a large fraction comprised low-complexity tandem repeats (MIC 359 Mbp, MAC 231 Mbp) (Figure S6). About one million interspersed repeats from 915 families were annotated in each genome assembly, covering 571 Mbp (MIC) and 454 Mbp (MAC), most of which were not assigned to a known repeat class (757 k copies, 366 Mbp total length in MAC). Interspersed repeat and low-complexity tandem repeat annotations overlapped substantially. Genome sizes after repeat masking were similar (MIC 245 Mbp, MAC 229 Mbp) and closer to k-mer based size predictions (Table S1). The difference in total assembly sizes is likely caused by misassembly of low-complexity repeats, rather than by imprecise elimination of repetitive elements in addition to precise IES excision, as found in the ciliate *Paramecium*,^44^ because the proportion of low complexity sequences in unassembled reads hardly differs between MIC and MAC (Figure 2A, Figure S7).

### Gene prediction for context-dependent sense/stop codons

Karyorelicts including *Loxodes* generally use an ambiguous stop/sense genetic code (NCBI translation table 27) where the only stop codon, UGA, can also encode tryptophan (W) if sufficiently far upstream of the mRNA poly(A) tail.^45,46^ Coding UGAs must be distinguished from stop UGAs to predict genes, but existing software does not permit single codons with alternative, context-dependent translation outcomes.

Assembled transcripts with poly-A tails ≥7 bp and with BLASTX hits to published ciliate proteins revealed informative sequence characteristics for predicting stop UGAs. Like other ciliates, 3’-untranslated regions (3’-UTRs) of *Loxodes* were relatively short (mean 53 bp, median 41 bp) (Figure 3A). Coding sequences (CDSs) were more GC-rich than 3’-UTRs (33.5% GC vs. 18.6% respectively), and showed a 3-base periodicity in their base composition associated with codon triplets (Figure 3B). Coding UGAs and UAAs were depleted for about 20 codon positions before the putative true stop UGA (Figure 3C), unlike other codons (Figure S8).

**Figure 3.**
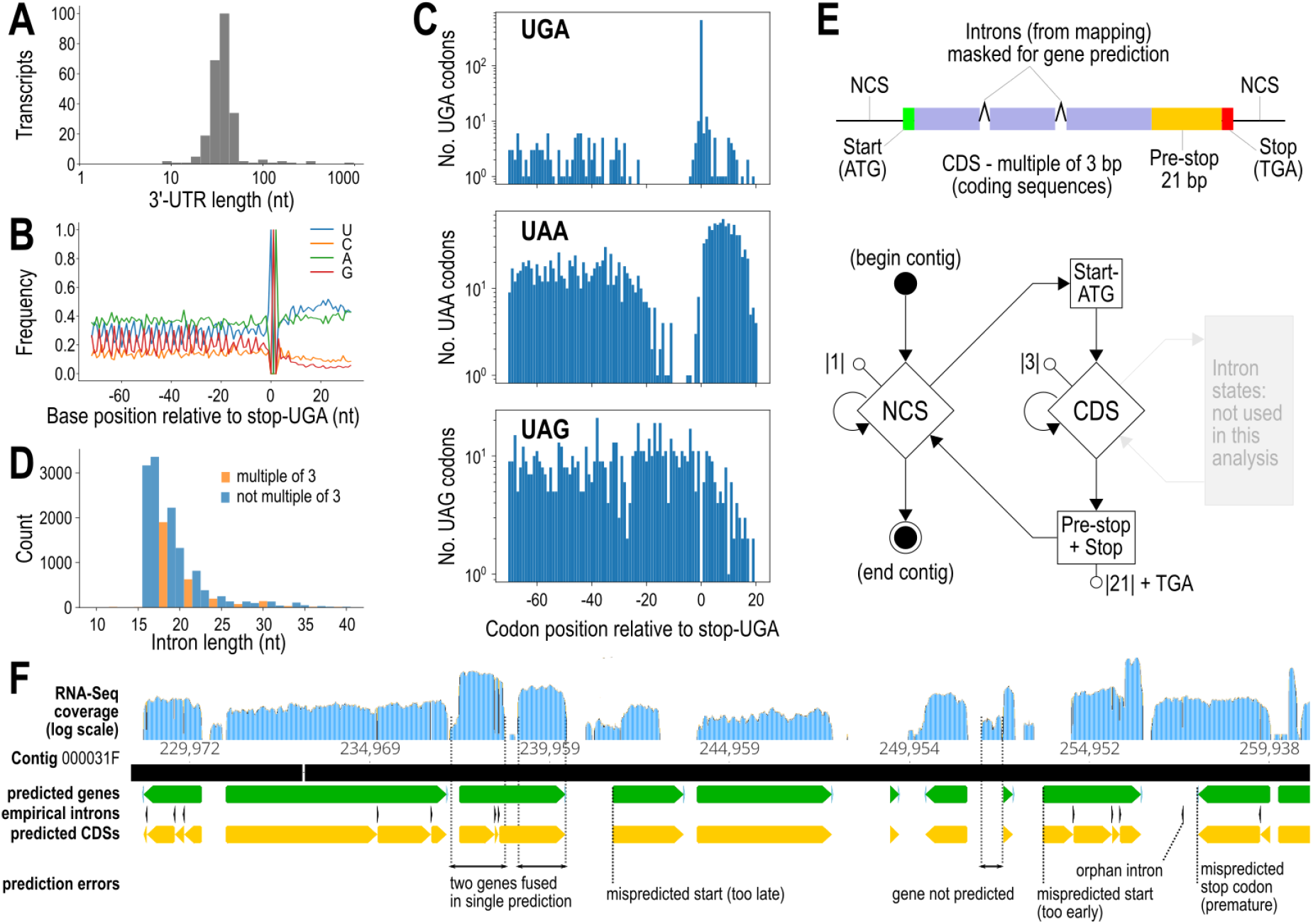
*Loxodes magnus* gene prediction. (A) Length distribution of 3’-untranslated regions (3’-UTRs) from poly-A tailed transcripts with stop codons predicted from BLASTX hits to other ciliates. **(B)** Base composition around predicted stop codons in transcripts. **(C)** Counts of UGA, UAA, and UAG codons relative to predicted stop-UGA codons in transcripts, showing depletion of in-frame UGA and UAA immediately upstream of stop-UGAs, but no depletion of UAG (see also Figure S7). **(D)** Length distribution of introns predicted from RNA-seq mapping to MAC genome assembly (excluding orphan introns). **(E)** Diagrams of gene model and GHMM used for gene prediction. Corresponding reverse-complement states (mirror image) are not shown. Introns were annotated empirically from RNA-seq mapping. **(F)** Excerpt of Pogigwasc gene prediction from MAC assembly contig 000031F, showing annotation tracks for predicted genes (green), CDSs (yellow), empirical introns (black), aligned against RNA-seq coverage (blue). Common types of mispredictions recognizable by comparison with RNA-seq mappings are indicated.

*Loxodes* introns identified by RNA-seq mapping to the MAC assembly were much shorter than in typical eukaryotes (mean 19.3 bp, mode 17 bp, 93% with length ≤25 bp; introns <16 bp appeared negligible and may be errors or outliers) (Figure 3D), but nonetheless longer than in heterotrichs, the sister group to karyorelicts, where almost all introns were 15 bp (∼95%) or 16 bp.^12,15^ Introns with lengths of a multiple of three (3*n*-introns) were relatively depleted (Figure 3D), as previously observed in oligohymenophorean and spirotrich ciliates.^47,48^

We designed a generalized hidden Markov model (GHMM) for *Loxodes* gene prediction with a probabilistic state for the codon UGA (either W or Stop), adapted from the model used by AUGUSTUS.^49^ Additionally, the stop UGA is preceded by a “pre-stop” region of 21 nt wherein no in-frame UGAs are permitted, to model the observed depletion of coding UGAs immediately upstream of the stop UGA (Figure 3E). *Loxodes* introns were difficult to model because of their short length and unusual length distribution, so we annotated them empirically from RNA-Seq mappings. We implemented the GHMM in our software Pogigwasc (https://github.com/Swart-lab/pogigwasc) and parameterized with a set of 152 manually annotated genes^50^ 94% completeness was estimated by BUSCO (Alveolata marker set) from the predicted proteome (Figure S9, Supplementary Results 4).

### Searches for genes and small RNAs related to genome editing

The *Loxodes magnus* genome assembly encoded no detectable homologs of proposed domesticated ciliate IES excisases. Neither the DDE_Tnp_1_7 (Pfam PF13843) domain found in PiggyBac family homologs (PiggyMacs, Pgm) of oligohymenophoreans and heterotrichs, nor the DDE_3 (PF13358) domain from TBE element transposases of the ciliate *Oxytricha* was annotated in *L. magnus* predicted proteins (Figure 4A). To account for incompletely predicted genes, we performed a translated search (TBLASTN) against the genomes with model ciliate Pgm and TBE-transposase protein sequences. The best hit (*Blepharisma stoltei* Pgm ^9^ to the *L. magnus* MIC) had an E-value of only 0.12, compared to 10^-33^ for an alignment of *Paramecium tetraurelia* Pgm to the *B. stoltei* genome that recovered the *B. stoltei* Pgm. The weak match in *L. magnus* is hence likely spurious.

**Figure 4.**
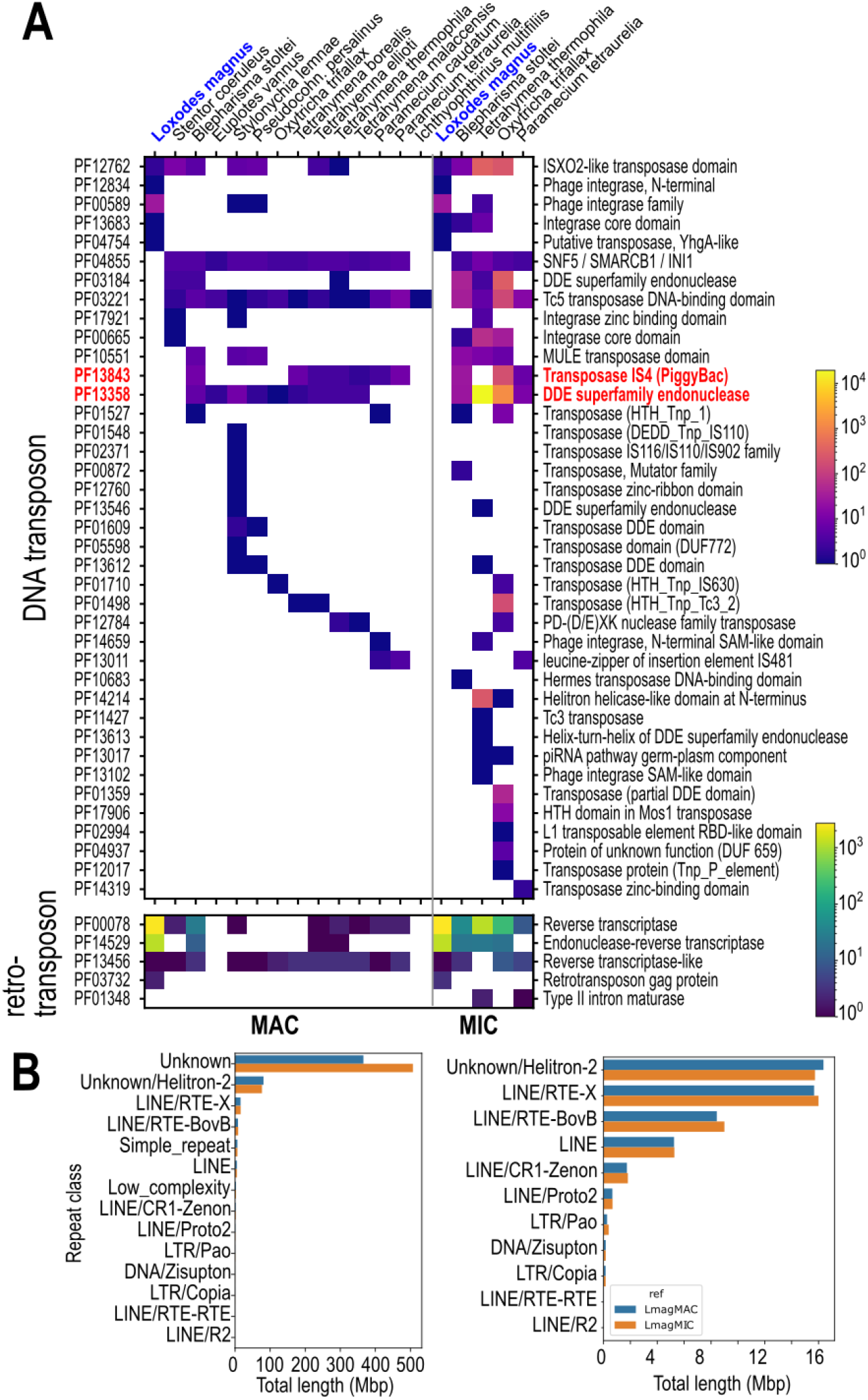
**Repeats and mobile element domains in *Loxodes* compared to other ciliates**. **(A)** Heatmaps representing Pfam domain counts related to mobile elements per MAC (left) or MIC (right) genome in ciliates. Red text: domains associated with known or proposed genome editing excisases (PF13843, PF13358). **(B)** Total lengths of interspersed repeat annotations in MIC vs. MAC genomes, sorted by classification. Left: All categories included. Right: “Unknown” and rnd-1_family-2 (“Unknown/Helitron-2”) excluded to show details.

Apart from the domesticated excisases, other components of the ciliate genome editing toolkit are difficult to distinguish from homologs with other functions. An exception are Dicer ribonucleases: ciliates have two Dicer classes: canonical Dicer (Dcr) for siRNA biogenesis, and development-specific Dicer-like proteins (Dcl) that lack additional Dcr N-terminal domains, which produce precursors to sRNAs involved in genome editing.^25,51^ Both Dcr and Dcl homologs were found in *L. magnus* (Figure S10, Supplementary Results 5).

We found no evidence for editing-associated small RNAs in *L. magnus*. If present, we reasoned that they should be produced in actively growing populations of *Loxodes* during asexual division where MIC-to-MAC development is obligatory, but not in starved populations without active division. However, sRNA length distributions in both actively growing and starved cells were similar (peaks 24, 25 nt). Development-specific sRNAs should map to both DNA strands, but the *Loxodes* sRNAs observed are strand-biased and probably represent antisense, gene-silencing siRNAs (Figure S11, Supplementary Results 6).

### Abundant retrotransposon-related vs. rare DNA transposon-related elements in Loxodes magnus

Thousands of copies of the retrotransposon-related domains reverse transcriptase RVT_1 (PF00078, ∼2700 copies) and endonuclease Exo_endo_phos_2 (PF14529, ∼1200 copies) were encoded in both nuclear genomes of *L. magnus*. This was ∼100 times the next highest counts in ciliates in the *Blepharisma stoltei* MAC genome,^12^ and contrasted with the paucity of DNA transposase-related domains (Figure 4A).

At least two repeat families, rnd-1_family-27 and rnd-1_family-19, appeared to represent complete long interspersed nuclear elements related to LINEs and other autonomous non-LTR retrotransposons with 5-6 kbp consensus length; only about 10% of the ∼3000 copies detected per family were full-length with low (<10%) sequence divergence from the consensus (Table S2). They contained coding sequences with both RVT_1 and Exo_endo_phos_2 domains typical of LINEs.^52^ The top BLASTp hits to GenBank’s nr database for representative *Loxodes* proteins encoding these domains were to *Blepharisma stoltei* proteins, so these elements may date to the karyorelict/heterotrich common ancestor. In total >30,000 repeat elements per genome assembly were classified by RepeatMasker as LINEs (Figure 4B), most of which were incomplete and hence likely inactive (Table S2). 504 instances of interspersed repeats overlapped closely with indel polymorphisms (>90% reciprocal overlap), including ten full-length copies of rnd-1_family-27 and two of rnd-1_family-19. The indels also help to confirm that mobile element family boundaries were correctly predicted, which is otherwise difficult for non-LTR retrotransposons because they may not be bound by conserved motifs or target site duplications.^53^

Unlike the retrotransposon sequences, repeats classified as helitrons or DNA transposons lacked the expected conserved domains and were likely to be spurious annotations (Supplementary Results 8). Additionally, two proteins with the “ISX02-like transposase” motif (PF12762, DDE_Tnp_IS1595) were related to sequences from *Blepharisma* and *Stentor* but probably no longer involved in transposition (Supplementary Results 9), and the gene containing a YhG-like transposase domain (PF04654) was associated with a gene cluster with signs of recent horizontal gene transfer from *Rickettsia* bacteria (Supplementary Results 10).

### *Loxodes magnus* MACs have characteristics of both active chromatin and heterochromatin

*L. magnus* nuclei have distinct morphology (Figure 1C) and chromatin organization. MAC protein composition was more diverse, as silver-stained PAGE gels revealed multiple prominent bands for MACs compared to few visible bands for MICs, of which the most prominent corresponded to typical histone sizes (Figure 5A).

**Figure 5.**
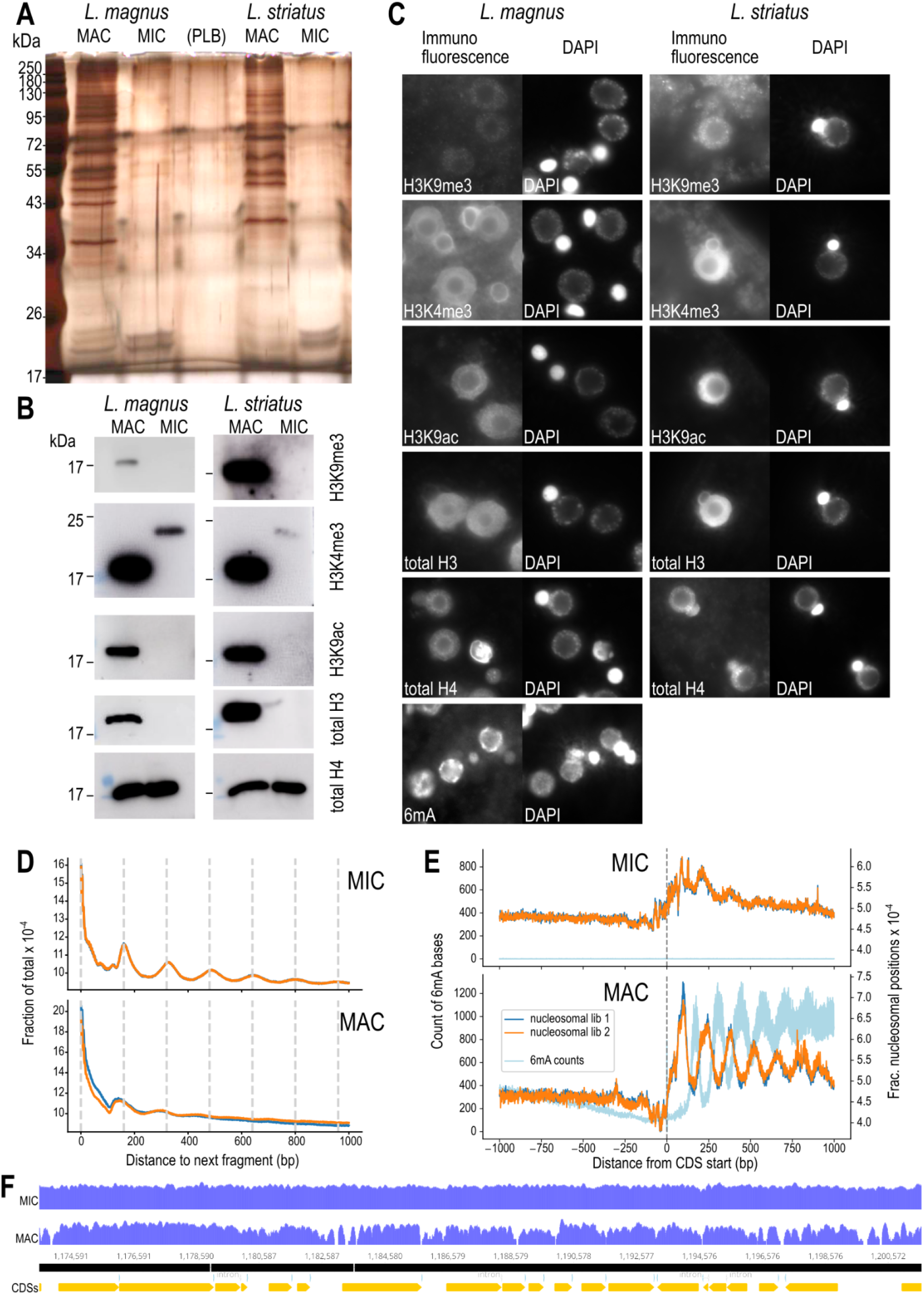
Molecular differences between *Loxodes* MIC and MAC nuclei. (A) Silver staining of protein extracts from flow-sorted MIC and MAC. PLB – protein loading buffer only. **(B)** Western blots against histone modifications in flow-sorted MIC and MAC. **(C)** Secondary immunofluorescence against histone modifications or 6mA, alongside DAPI staining of DNA. Panel widths: 30 µm. **(D)** Global phaseograms of nucleosomal DNA density (two replicates: dark blue, orange lines) in flow-sorted MIC and MAC for *L. magnus* (low complexity repeats masked); vertical lines – 160 bp intervals. **(E)** Phaseograms of nucleosomal DNA density (two replicates: dark blue, orange lines) and 6mA modified bases (light blue) relative to predicted coding sequence start positions in flow-sorted MIC and MAC of *L. magnus*. **(F)** Example of coverage pileups (log scaled) for MIC vs. MAC nucleosomal DNA reads mapped to MAC reference assembly (contig 000000F), aligned with CDS predictions (bottom track).

Histone marks typical of both activation and repression were detected by Western blots in MACs but not in MICs (Figure 5B), namely histone H3 lysine 9 acetylation (H3K9ac, active transcription) and H3 lysine 9 trimethylation (H3K9me3, heterochromatin). H3 lysine 4 trimethylation (H3K4me3, euchromatin) was detected in MACs at the expected size (∼17 kDa) but MICs showed a weaker, higher-weight band. Immunofluorescence localization was consistent with Western blots (Figure 5C). As expected, histone marks in MACs colocalized with DAPI-stained chromatin but were absent from nucleoli, although signals for H3K9me3 and H3K4me3 had background signal in the cytoplasm, and H3K4me3 also showed a peripheral signal surrounding MICs that was not colocalized with DNA.

Total histone H3 was detected in MACs but not MICs with a commercial antibody (Figure 5B, 5C). The *Loxodes magnus* genome encodes multiple histone H3 homologs, clustering into three groups, only one of which (canonical H3-related) was likely to be detected by the antibody applied (Figure S12, Supplementary Results 11), hence MACs likely use canonical H3 while MICs may use a different variant. In contrast, histone H4, the most conserved core histone, was detected in both nuclei (Figure 5B, 5C).

Nucleosomal positioning patterns differed between MIC and MAC at both the global scale and relative to gene features. Similar dsDNase digestion conditions to isolate nucleosomal DNA yielded smaller fragments from MACs than MICs (Figure S13A). When sequenced and mapped to the genome, the global phaseogram, i.e. the distribution of nucleosomal fragment positions relative to each other, displayed periodic peaks at multiples of 160 bp, the expected length of nucleosomal plus linker DNA (Figure 5D). These peaks were more pronounced from MICs than MACs. In contrast, phaseograms relative to the starts of predicted coding sequences showed periodic peaks within coding sequences in the MAC but not in the MIC (Figure 5E). Coverage pileups of MAC nucleosomal reads also showed arrays relative to gene features, which were not seen with MIC nucleosomal reads (Figure 5F). We interpret this to mean that MIC chromatin is condensed and largely inactive, with nucleosomes regularly arrayed but positioned independently of gene locations, whereas the MAC has more accessible DNA where nucleosomes are arrayed relative to genes due to transcription (Figure S13C).

Though cytosine methylation has been reported in some ciliate species, it is apparently absent in others, and no canonical cytosine DNA methyltransferases have been identified yet.^33,54,55^ We were also unable to detect such methyltransferases in *Blepharisma*. In contrast, the base modification 6mA was abundant and found predominantly in the *Loxodes* MAC by both immunofluorescence (Figure 5C) and PacBio SMRT-Seq (4,405,028 ApT positions (0.85%) in MAC vs. 845 (0.00013%) in MIC genome). 99.6% of base modification calls were in ApT motifs, which are also the exclusive motif for 6mA in *Tetrahymena*.^56^ 6mA across *Loxodes* gene bodies showed a distinct periodicity with alternate phase to the nucleosome positioning (Figure 5E), similar to *Tetrahymena* and *Oxytricha*.^34,36^ Unlike in other ciliates, 6mA coverage does not fall off sharply towards the 3’ end of the gene body. 6mA was largely absent from *Loxodes* genes not transcribed by RNA polymerase II (e.g. rRNA), suggesting that 6mA methylation is coupled to RNA polymerase II transcription, like in *Tetrahymena*.^36^

Almost all ApT motifs with 6mA in *Loxodes magnus* were hemi-methylated in both MAC (99.87%) and MIC (100%) assemblies (e.g. Figure S11C), unlike other ciliates, which have a mixture of 6mA hemi- and full methylation in MACs, including *Blepharisma* (59.4% hemi), *Tetrahymena* (11% hemi),^56^ and *Oxytricha*.^34,36^ Full methylation is necessary for semi-conservative 6mA transmission during asexual MAC division.^56^ Therefore absence of full methylation in *Loxodes* is consistent with their non-dividing MACs and with *de novo*, non-epigenetic methylation during MAC formation. The *Loxodes* genome also lacks homologs of the *Tetrahymena*/*Oxytricha* 6mA methyltransferase complex p1 and p2 components, suggesting they are not needed for hemi-methylation (Supplementary Results 12).

## Discussion

The karyorelictean ciliate *Loxodes magnus* maintains morphologically, molecularly, and functionally distinct somatic vs. germline nuclei, but in a counterpoint to other ciliates, we did not detect extensive genome editing in the form of IES excision or differential amplification.

### Comparison with previous studies

Our conclusions contradict a previous report of genome editing in an uncultivated *Loxodes* sp.^43^ Their claim of up to 10^4^-fold variation in genome amplification is quantitatively unrealistic and is likely a methodological artifact, whereas their putative IESs resemble the indel polymorphisms observed in this study (Supplementary Discussion). Nonetheless, other karyorelicts may have some degree of editing: developing MACs of *Trachelonema sulcata* have distinctly less DNA than MICs or mature MACs, suggesting some DNA elimination in early MAC development.^57^ Old MACs in *Loxodes* have higher and more variable DNA content than recently matured MACs,^38^ but this may be nonspecific amplification in senescent nuclei, as we did not observe distinct subclusters of MACs by DNA content (Figure 1D) nor evidence for differential amplification. Chromosome breakage and unscrambling were not directly addressed here, but unscrambling has only been found in conjunction with IES elimination,^5,58^ and is likely also absent. To assess chromosome breakage, *Loxodes* telomeres, which have thus far eluded our detection, will need to be identified.

### Implications for mobile element proliferation and management

The large *Loxodes* MAC genome with more mobile elements and repeats than typical ciliates is consistent with the expected evolutionary consequences of losing or drastically reducing IES excision, but not because genome editing was a “defense” against mobile elements, as it has often been characterized.^59,60^ It has recently been argued that editing actually helps mobile elements persist in the germline by shielding them from selection in the soma, and that editing is maintained by evolutionary addiction rather than positive selection.^8,9,28^ *Loxodes* supports our interpretation by showing that editing is not necessary for survival per se. How, then, does it ameliorate the deleterious effects of mobile elements?

Natural selection would eliminate the most deleterious elements, because they are “exposed” in the transcriptionally active MAC genome in *Loxodes*. For instance, repeats that correlate with indel polymorphisms have not yet reached fixation and are hence liable to selection. Those that remain must be largely inactive or benign, such as the abundant but mostly fragmentary retrotransposon-related repeats (retrotransposons are known to be prone to incomplete reverse transcription that results in truncated fragments.)^61^ Though retrotransposons outnumber DNA transposons in many eukaryotes, e.g. 43% vs. 4% of the human genome,^62^ it is surprising that we have not detected compelling DNA transposon homologs in *Loxodes*, as they are numerous in all ciliate MIC genomes examined to date (Figure 4A).^5,6,8,9,58^ This may simply reflect the most recent wave of mobile element proliferation in this particular strain, or higher deleteriousness of DNA transposons than retrotransposons.

*Loxodes* may also have maintained ancestral mechanisms that suppress mobile element expression that other ciliates have extended to editing. Morphologically mature *Loxodes* MACs have the heterochromatin-associated histone mark H3K9me3 (Figure 5C), whereas other ciliates (with genome editing) have H3K9me3 only in developing MACs but not mature MACs or MICs, because they have co-opted heterochromatin histone marks H3K9me3 and H3K27me3 to guide the editing machinery,^63,64^ although low H3K27me3 levels have been reported in *Paramecium* mature MACs.^65^

### Secondary loss or retention of ancestral state?

Evolutionarily, either IES excision was present in the ciliate last common ancestor (LCA) but secondarily lost in Karyorelictea, or it was absent in the ciliate LCA and karyorelicts reflect the ancestral state. Secondary loss is more parsimonious, because karyorelicts are sister to heterotrichs,^42,66^ in which at least one genus (*Blepharisma*) performs extensive genome editing like other ciliates.^9,12^ The presence of Dcl genes in *Loxodes*, homologous to those involved in genome editing in other ciliates, also support secondary loss, whereas the apparent absence of a domesticated excisase is less conclusive, as ciliate excisases come from at least two different families,^12,19,21,67^ and so were independently or repeatedly domesticated.

### Possible scenarios for loss of IES excision

In the evolutionary addiction model, ciliates with many intragenic IESs like *Paramecium* or *Blepharisma* cannot afford to lose genome editing as the resulting erroneous retention of IESs in essential genes is likely lethal. Conversely, IESs cannot be exposed to selection in the somatic genome if they are removed by editing. How then can IESs or genome editing be lost? We see three possibilities for how the ancestor of *Loxodes* could have lost genome editing in the face of this conundrum: (i) its IESs were mostly intergenic and nonlethal if retained; (ii) high rates of gene duplication such that some paralogs remained undisrupted by IESs; or (iii) a mature MAC without IESs was developmentally “reset” to a MIC, wiping the germline clean of IESs in one go.

The loss of asexual MAC division likely preceded loss of genome editing in karyorelicts, as the increased cost of additional MIC-to-MAC development during asexual division could cause strong selective pressure to streamline or lose genome editing. In other ciliates, development of MIC precursors into MACs is coupled to the sexual cycle, and MIC-to-MAC development is costly compared to asexual division because of genome editing, e.g.

*Paramecium* requires ∼22 h for sexual vs. 6 h for asexual division.^68,69^ Asexual MIC-to-MAC development without prior meiosis/karyogamy has actually been observed in *Blepharisma*, where “somato-MICs” develop into MACs if developing MACs are experimentally removed,^70^ although genome editing is presumably still involved since its MIC genome possesses ∼40,000 IESs. Karyorelict MIC-to-MAC development may be developmentally homologous to this “backup” somato-MIC pathway.

### Genome editing may not be essential to nuclear dualism

Even without genome editing, nuclear dualism per se may hinder mobile element invasion because MIC chromatin is condensed and inactive for most of the life cycle, whereas any successful invasion of the disposable somatic MAC would not be transmitted to progeny, both sexual and asexual in the case of *Loxodes*. An inactive MIC would also limit transcription-associated mutation, thereby maintaining germline DNA integrity.

Though many challenges remain in imagining how exactly a cell goes from one to two functionally differentiated types of nuclei, *Loxodes* suggests that a prototypical characteristic of model ciliates, extensive genome editing, is not obligatory. Broader taxonomic sampling of both MIC and MAC genomes will be needed to ascertain if all ciliates with dividing MACs also have genome editing, and conversely if all karyorelict ciliates which have non-dividing MACs also appear to lack genome editing.

## Materials and Methods

General reagents were analytical grade and purchased from Sigma-Aldrich/Merck unless otherwise noted. Full parameters of computational analyses are available from code repositories linked below. R.T. – room temperature.

### Isolation and cultivation of Loxodes strains

Strains *Loxodes magnus* Lm5 and *Loxodes striatus* Lb1 were isolated from single cells and grown in soil extract medium as previously described.^73^

### Nuclei purification by fluorescence-activated nuclear sorting

500 mL batches of dense culture (∼500 cells/mL) were starved for at least the average doubling time of ∼1 week,^73^ filtered through pre-washed quartz sand, centrifuged (120 g; 2 min; R.T.) in pear-shaped glass flasks, resuspended in autoclaved Volvic water, concentrated by centrifugation to ∼3 mL, then resuspended in 7.5 mL ice-cold lysis buffer (sucrose 0.25 M, MgCl_2_ 10 mM, Tris-HCl pH 6.8 10 mM, Nonidet P-40 0.2% w/v)^74^ in 15 mL polypropylene tubes. The mixture (on ice) was pulled up and expelled completely five times with a 20 mL plastic syringe through a 0.60 mm × 60 mm needle to lyse cells, stained with DAPI (final conc. 1 µg/mL), transferred to 2 mL tubes, and centrifuged (2000 g; 3 min; 4 °C); supernatant was removed by pipetting, and the nucleus pellet resuspended in 2 mL ice-cold Galbraith’s buffer (MgCl_2_ 45 mM, sodium citrate 30 mM, MOPS pH 7 20 mM, Triton X-100 0.1% v/v)^75^ by pipetting up and down, then kept on ice until sorting.

Suspensions were filtered through 35 µm nylon mesh “cell strainers” (Fisher Scientific 352235), then sorted on a BD FACSMelody Cell Sorter, controlled with BD FACSChorus v1.1.18.0, with 100 µm nozzle size, 23 PSI pressure, 34.0 kHz drop frequency, and “purity” sort mode. DAPI fluorescence was measured with 405 nm laser excitation and 448/45 filter.

PMT voltages were set to initial values: FSC, 300 V; DAPI, 370 V; SSC, 490 V. Populations were gated with combinations of SSC, FSC, and DAPI fluorescence (Figure 1D), but exact settings were adjusted manually to account for batch variation.

Sorted nuclei were collected in 1.5 mL microcentrifuge tubes pre-filled with 100 µL Galbraith’s buffer, cooled to 5 °C. A 10 µL sample of each batch of sorted nuclei was viewed under epifluorescence microscopy (DAPI signal) to verify sorting purity. At least 100 nuclei per sample were counted and scored as MIC (no nucleolus) or MAC (with nucleolus). Only samples with >99% visually verified purity of the target nucleus type were used for downstream experiments. Collected nuclei were centrifuged (8000 g; 3 min; 4 °C), supernatant was removed by pipetting; pellets were then snap-frozen in liquid nitrogen and stored at -80 °C until use.

Detailed protocol: https://doi.org/10.17617/3.OYUXDS.

### Genomic DNA library preparation and sequencing

DNA was isolated from sorted nuclei with the CleanNA Clean Blood and Tissue kit (CBT-D0096), resuspended in 10 mM Tris-HCl pH 8.5, and quantified with the Qubit DNA High Sensitivity kit. Short-read libraries were prepared with the NEBNext Ultra II FS DNA Library Prep kit (NEB E7805), and sequenced 150 bp paired-end on an Illumina HiSeq3000. Long-read libraries were prepared with the SMRTbell Express Template Prep Kit 2.0 using the Sequel II Binding Kit 2.0 with Sequel polymerase 2.0, size-selected to 10 kbp, and sequenced with the HiFi protocol on a PacBio Sequel II.

### RNA library preparation and sequencing

*Loxodes magnus* cells grown in soil extract medium,^73^ were resuspended in fresh medium to target density of 250 cells/mL, split into six flasks of 150 mL each, and kept at R.T. without feeding. Cell densities were monitored daily by counting cells in 3×100 µL aliquots per flask (Table S3). Three flasks representing “starved” cells were harvested for RNA extraction after three days. The remaining flasks were each fed with 450 µL of concentrated *Chlamydomonas* ^73^ on days 3 and 4. By day 5, dividing *Loxodes* cells were observed and cell densities began to recover, so flasks were harvested, representing “fed” cells.

To harvest, cells were filtered through cotton gauze, centrifuged in pear-shaped flasks (80 g; 1 min; R.T.), resuspended in 10 mL SMB medium in 15 mL polypropylene tubes, and centrifuged (90 g; 1 min; R.T.). Concentrated cells (∼500 µL) were transferred dropwise to 3 mL ice-cold TRI reagent (Sigma-Aldrich T9424) while vortexing, and stored at -80 °C until use. For RNA extraction, thawed samples were split into 3×1 mL aliquots. Each aliquot was shaken with 200 µL chloroform, kept at R.T. for 2 min, then centrifuged (1200 g; 15 min; 4 °C). The aqueous phase was transferred to new tubes, mixed with equal volume 100% ethanol, inverted 20×, then purified with Zymo RNA Clean and Concentrator 5 kit (Zymo, R1013) with in-column DNase digestion.

### Nucleosomal DNA library preparation and sequencing

*Loxodes magnus* cells were harvested and washed once as described above, then centrifuged (200 g; 1 min; R.T.). The cell pellet was resuspended with ice-cold Galbraith’s buffer amended with bovine serum albumin (BSA, 0.05% w/v) and cOmplete protease inhibitor (1×, Roche 11697498001), lysed by repeated pipetting, stained with DAPI (1 µg/mL) for 5 min on ice, centrifuged (500 g; 2 min; 4 °C), and resuspended again in Galbraith’s + BSA + protease inhibitor on ice. Nuclei were flow-sorted as described above.

Nucleosomal DNA was prepared with the EZ Nucleosomal DNA prep kit (Zymo D5220). Sorted nuclei were centrifuged (1000 g; 2 min; 4 °C), and supernatant removed by pipetting. Each nuclei pellet was washed with 100 µL ice-cold Atlantis digestion buffer, centrifuged (200 g; 1 min; 4 °C), resuspended in cold 50 µL Atlantis digestion buffer by 10× repeated pipetting, then incubated with 10 units Atlantis dsDNase (42 °C; 40 min). Stop solution was added, then DNA was purified on spin columns and eluted in 30 µL buffer. Mono- and di-nucleosomal DNA fragments (∼150 to 300 bp) were size-selected with SPRIselect magnetic beads (Beckman-Coulter B23317) using “right size selection” and 0.7× beads:sample volume ratio. Fragments were sized with a Bioanalyzer 2100 DNA high sensitivity assay (Agilent 5067-4626). Libraries were prepared with the NEBNext Ultra II DNA library prep for Illumina kit (NEB E7645S) and sequenced on an Illumina NextSeq2000.

### Nucleosomal DNA profiling and phaseograms

Nucleosomal DNA libraries for MAC and MIC were mapped onto the MIC Falcon reference assembly with minimap2 v2.24 with parameter: -ax sr. Positional maps, or “phaseograms”, were computed with mnutils commit 105d129 (https://github.com/Swart-lab/mnutils), with parameters: --feature gene --phaseogram --dump, using gene features predicted by Pogigwasc in GFF3 format. The insert size range (represented by parameters --min_tlen and --max_tlen) were set to 96-136 bp for MAC and 126-166 bp for MIC, because nucleosomal DNA was more heavily digested in MAC than MIC. Read mappings without peak-calling or denoising were used to obtain a purely empirical picture of nucleosomal positioning. For all phaseograms, the midpoint of each mapped read pair was used as the nucleosomal DNA fragment position. For each mapped fragment, positions of other fragments in a 1 kbp window downstream were enumerated; the cumulative pileup of positions relative to each other constituted the global phaseogram. The Pogigwasc gene predictor only modeled coding sequences. Therefore, for the phaseogram relative to gene features, we assumed that 5’-UTR lengths are short and tightly distributed like other ciliates, and used coding sequence starts as a proxy for transcription start sites, using a window of 1 kbp on both sides.

Workflow: https://github.com/Swart-lab/loxodes-nucleosomes-workflow

### k-mer based genomic library comparisons

Adapter- and quality-trimmed (Phred score >28) Illumina reads were used for k-mer based comparisons. k-mer content (k=21) of genomic libraries were compared pairwise with each other, or with the reference MAC genome, using the ‘kat comp’ command in kat v2.4.2,^76^ which depends on jellyfish^77^ and SeqAn.^78^

Workflow: https://github.com/Swart-lab/loxodes-kmer-comp

### Genome assembly

PacBio sequencing reads were demultiplexed and processed to circular consensus sequence (CCS) reads with PacBio SMRT Link v9. CCS reads were first assembled with Flye v2.8.1 ^79^ using the option: --pacbio-hifi. A preliminary analysis showed that the genome was likely to be diploid, therefore CCS reads were assembled again with the diploid-aware assembler Falcon (Bioconda package pb-falcon 2.2.4 installed with package pb-assembly v0.0.8)^80^ using a relatively low identity threshold of 0.96 for collapsing heterozygosity (option: overlap_filtering_setting = --min-idt 96) and option: ovlp_daligner_option = -e.96. Other parameters followed the suggested configuration template for CCS reads (https://github.com/PacificBiosciences/pb-assembly/blob/master/cfgs/fc_run_HiFi.cfg). The average coverage (∼20-30x) was below the recommended ∼30x coverage per haplotype for phased assembly, so we did not proceed to Falcon-Unzip. Falcon primary contigs were polished with Racon v1.4.20 ^81^ using read mappings from pbmm2 v1.4.0 filtered with samtools view using options -F 1796 -q 20 (exclude records for unmapped reads, non-primary alignments, reads that fail platform/quality checks, and PCR or optical duplicates; minimum quality Phred 20).

Workflow: https://github.com/Swart-lab/loxodes-assembly-workflow

### Annotation of repeats in genome assembly

Low-complexity tandem repeats were annotated with TRF v4.09.1 ^82^, using the recommended algorithm settings: 2 5 7 80 10 50 2000 -d -h -ngs. The output was filtered and converted to GFF format with trf_utils (https://github.com/Swart-lab/trf_utils), retaining repeat regions ≥ 1 kbp long; if features overlapped, the highest-scoring feature was retained, otherwise the feature with the most repeat copies. The filtered feature table was merged and used to mask the assembly with the merge and maskfasta commands in bedtools v2.27.1.^83^

Interspersed repeat element families were predicted from the MIC genome assembly with RepeatModeler v2.0.1 (default settings, random number seed 12345) with the following dependencies: rmblast v2.10.0+ (http://www.repeatmasker.org/RMBlast.html), TRF 4.09,^82^ RECON,^84^ RepeatScout 1.0.6,^85^ RepeatMasker v4.1.1 (http://www.repeatmasker.org/RMDownload.html). Repeat families were also classified in the pipeline by RepeatClassifier v2.0.1 through comparison against RepeatMasker’s repeat protein database and the Dfam database. Predicted repeat families were annotated in both the MAC and MIC assemblies with RepeatMasker, using rmblast as the search engine.

### Transcriptome mapping and assembly

RNA-seq libraries were adapter- and quality-trimmed (Phred > 28, length ≥ 25 bp) with bbduk.sh from BBtools v38.22. Reads were mapped with bbmap.sh (BBtools) to the *Chlamydomonas reinhardtii* reference genome (JGI Phytozome assembly v5.0, annotation v5.6)^86^ (identity ≥0.98) to remove potential contamination from food algae. RNA-seq reads were mapped to reference genome assemblies with Hisat2 v2.0.0-beta,^87^ modified to lower the minimum allowed intron length to 10, with options: --min-intronlen 10 --max-intronlen 50000 --seed 12345 --rna-strandness RF.

Workflow: https://github.com/Swart-lab/loxodes-assembly-workflow

### IES prediction

PacBio CCS reads were mapped to the reference Falcon assembly with minimap2 v2.17 ^88^ with the options: --MD -ax asm20. BAM files were sorted and indexed with samtools v1.11.^89^ Putative IESs were predicted from the mapping BAM file with BleTIES MILRAA v0.1.11 ^90^ in CCS mode with options: --min_break_coverage 3 --min_del_coverage 5 --fuzzy_ies --type ccs, parallelized with ParaFly commit 44487e0 (https://github.com/ParaFly/ParaFly).

Workflow: https://github.com/Swart-lab/loxodes-bleties-workflow

### Variant calling and comparison to putative IESs

Variants were called with Illumina short reads (more accurate, higher coverage), whereas phasing and haplotagging were performed with PacBio long reads, as recommended in the WhatsHap documentation. Illumina MIC and MAC reads were mapped to MAC reference assembly with bowtie2 v2.3.5 ^91^ with default parameters. Variants were first called from mapped Illumina reads with FreeBayes v1.3.2-dirty ^92^ in “naive” mode to verify ploidy, with options: -g 400 --haplotype-length 0 --min-alternate-count 1 --min-alternate-fraction 0 --pooled-continuous, filtered with vcffilter from vcflib v1.0.0_rc2 ^93^ to retain variant calls with Phred quality score > 20. Variants were then called again in diploid mode, i.e. with default options except: -g 400. PacBio HiFi reads mapped to the MIC and MAC assemblies were phased and haplotagged with WhatsHap v1.4,^94^ using only SNPs (default). VCF files were processed (e.g. merging, indexing) with bcftools v1.11.^89^ Reads with/without “IES” indels predicted by BleTIES were compared with their respective haplotags by parsing the haplotagged reads. The script used the pybedtools ^83,95^ and pysam ^96^ libraries.

Workflow: https://github.com/Swart-lab/loxodes-bleties-workflow

### Generalized hidden Markov model (GHMM) for gene prediction with ambiguous genetic codes

Published gene prediction software tools assume that stop codons are deterministic. We therefore modified a generalized hidden Markov model (GHMM) for eukaryotic genes ^49^ to accommodate ambiguous stop codons, implemented in the Java software package Pogigwasc (https://github.com/Swart-lab/pogigwasc). Technical details of the model and implementation are described in ^50^. Briefly, genome sequence is modeled as a sequence of the following hidden states (Figure 3A): Upon initiation, the model enters non-coding sequence (NCS) state; NCS emits 1 nt, then either loops back to NCS, or enters forward-strand Start or reverse-strand Stop states; genes can be encountered in either orientation, and are correspondingly represented by two sets of states. “Start” emits a 3 nt Kozak consensus sequence followed by a deterministic AUG start codon, then enters coding sequence (CDS) state. “CDS” emits 3 nt (one codon), then either loops back to CDS or enters “Stop” state. To avoid overfitting, codon emission probabilities follow a simplified model where the three codon positions are assumed to be independent, each drawing from the four possible nucleotide probabilities. “Stop” emits 24 nt, comprising 21 nt (seven codons) where the codon UGA is forbidden, followed by the UGA stop codon, then enters the NCS state. An intron model in the software was not used in this study.

To train and test the model, genes were manually annotated on the SPAdes preliminary assembly, based on alignments with assembled poly-A-tailed transcripts, mapping of RNA-Seq reads, and BLASTX alignments to other ciliate proteins. 152 genes were used for model training and 52 for testing.

### Gene prediction with Pogigwasc

Introns were empirically annotated from RNA-seq mappings using the Intronarrator pipeline commit b6abd3b (https://github.com/Swart-lab/Intronarrator), because their short length made it difficult to model them effectively, as previously observed with *Blepharisma*.^12^ Introns were identified from Hisat2 mappings of RNA-seq reads vs. the MAC and MIC Falcon assemblies with the following Intronarrator parameters: MIN_INTRON_RATIO=0.2, MIN_INTRONS=10, MAX_INTRON_LEN=40. Introns were then removed from the sequence to produce an artificial “intronless” assembly; non-coding RNAs were identified with Infernal v1.1.4 ^97^ and hard-masked. Scaffolds were split on Ns (scaffold gaps and hard-masked sequences) to produce pure contigs; contigs < 1 kbp were removed. Protein coding sequences were predicted from the resulting “intronless” contigs with Pogigwasc v0.1 with option: --no-introns, using the parameters trained on *Loxodes magnus*, which are bundled with the software. Annotations were translated back to the original genomic coordinates with scripts from pogigwasc-utils commit 7844e1 (https://github.com/Swart-lab/pogigwasc-utils). Gene predictions overlapping with low complexity regions predicted by TRF (see “Annotation of repeats in genome assembly”) were identified with bedtools intersect (options: -v -f 1.0).

Workflows: https://github.com/Swart-lab/loxodes-pogigwasc-workflow https://github.com/Swart-lab/loxodes-intronarrator-workflow

### Functional genome annotation and screening for genome editing toolkit

The *Loxodes magnus* predicted MIC and MAC proteomes from Pogigwasc, MAC proteomes from 13 ciliate species, and translated ORFs >30 a.a. predicted by getorf (EMBOSS v6.6.0.0) from MIC genomes of 4 species (Table S4), were annotated with InterProScan v5.57-90.0.^98^ Protein domains, signatures, and motifs relevant to the following functions were shortlisted by keyword searches of the InterPro database:^99^ DNA transposons and retrotransposons, Dicer and Dicer-like proteins, and histones (shortlists: https://doi.org/10.17617/3.BOFMWS). For retrotransposons, domains not relevant to mobile elements (e.g. telomerase reverse transcriptase) were excluded: Pfam domains PF00026, PF12009, PF11474. To account for the possibility that the coding sequence was not correctly annotated by Pogigwasc, domesticated excisases from the following ciliates were aligned against the *Loxodes magnus* genome assembly with TBLASTN (Blast+ v2.12.0)^100^ : PiggyMac homologs from *Paramecium tetraurelia* (Pgm, ParameciumDB PTET.51.1.P0490162), *Tetrahymena tetraurelia* (Tpb2p, Ciliate.org TTHERM_01107220), *Blepharisma stoltei* (BPgm, Ciliates.org BSTOLATCC_MAC17466), TBE element excisase from *Oxytricha trifallax* (Genbank AAB42034.1).

### Western blotting and immunofluorescence for histones and histone marks

Commercially available primary antibodies were used against the following histones or histone modifications: acetyl histone H3 lysine 9 (H3K9ac), trimethyl histone H3 lysine 9 (H3K9me3), trimethyl histone H3 lysine 4 (H3K4me3), total histone H3, and total histone H4 (Table S5). Western blotting with two additional antibodies was not successful: anti-trimethyl histone H3 lysine 27 (H3K27me3, Merck 07-449) (its 6 a.a. immunogen sequence was not found in *Loxodes* histone H3), and anti-histone H4 from Santa Cruz (sc-25260) (raised against human histone H4).

Nuclear pellets from flow sorting were resuspended with 1× protein loading buffer (PLB, 100 mM Tris-HCl pH 6.8, 4% (w/v) sodium dodecyl sulfate, 20% (w/v) glycerol, 0.2 M dithiothreitol, 0.05% (w/v) bromophenol blue) diluted with PBS (1000 nuclei per 1 µL final volume), and heated (95°C, 10 min). For each lane, 10 µL of sample in PLB was loaded onto a 12% SDS-PAGE gel and separated (200 V; 45 min) on a Bio-Rad Mini-Protean Tetra Cell electrophoresis system. Silver staining was performed with the Pierce Silver Stain Kit (24612, Thermo Fisher Scientific). For Western blots, proteins were transferred (80 V; 2 h; 4°C) onto a 0.2 µm nitrocellulose membrane (Bio-Rad 1620112). Membranes were air-dried, blocked with 5% (w/v) Bovine Serum Albumin (BSA) (Sigma A9647) with 0.2% (v/v)

Tween-20 (Sigma P2287) in PBS (overnight; 4°C), incubated with primary antibodies diluted in 5% BSA / 0.2% Tween-20 / PBS (overnight, R.T.), washed in 0.2% Tween-20 / PBS (3 × 10 min), incubated in the secondary antibody horseradish peroxidase (HRP)-conjugated goat anti-rabbit IgG (Merck 12-348) (1 h; R.T., washed with 0.2% Tween-20 / PBS (3 washes × 10 min), then washed in PBS (5 min). 200 µL of chemiluminescence substrate (Immobilon Crescendo Western HRP, Millipore, WBLUR0100) was added to each membrane, which was then imaged on a AI600 imager (GE Healthcare).

For Coomassie staining, 10 µL of resuspended protein samples in PLB were loaded on a 12% SDS-PAGE gel; samples were run in 1× Laemmli Buffer (Tris-base, Glycine, SDS) at 180 V until the loading dye ran out of the gel. The gel was stained with Coomassie blue (PhastGel blue R, Sigma, 6104-59-2) (overnight on orbital shaker; R.T., removed from staining solution, washed with autoclaved double distilled water (2 x 5 min), destained (25% v/v isopropanol, 10% v/v acetic acid in deionized water) until protein bands were clearly visible, then imaged with an AI600 imager.

For immunofluorescence, 100 mL of dense culture was centrifuged in pear-shaped flasks (80-120 g; 1 min; RT), resuspended in SMB medium to wash, centrifuged again, resuspended in 500 µL of SMB medium, and fixed at R.T. with an equal volume of ZFAE fixative.^73^ Subsequent transfers were performed by centrifugation (1000 g; 1 min) followed by removal of supernatant and resuspension of pellet at R.T. Fixed cells were permeabilized 5 min in 1.5 mL 1% (w/v) Triton-X / PHEM, post-fixed 10 min in 1 mL 2% (w/v) formaldehyde / PHEM, then washed twice for 5-15 min in 1 mL 3% (w/v) BSA / TBSTEM. Antibodies were diluted to working concentrations (Table S5) in 3% BSA / TBSTEM. The secondary antibody was Alexa Fluor 568-conjugated goat anti-rabbit IgG (Life Technologies, A11011). Fixed cells in BSA were incubated 10-60 min in primary antibody working solution, washed 5-10 min in 3% BSA / TBSTEM, then incubated 10-30 min in the secondary. Cells were counterstained ≥5 min with DAPI (1 µg/mL in 3% BSA / TBSTEM), mounted under ProLong Gold (Thermo Fisher), and cured (overnight; R.T.), then imaged by epifluorescence on a Zeiss AxioImager Z1 (Plan-Apochromat 63×/1.40 oil objective, Axiocam 702 camera, filter cubes Zeiss 49 for DAPI and AHF F46-008 for Alexa Fluor 568).

### 6mA base modification analysis from PacBio SMRT-Seq reads

PacBio SMRT-Seq subreads for flow sorted MAC and MIC DNA were indexed with pbindex (PacBio SMRT Link v12.0.0). Falcon assemblies were indexed with samtools faidx.

Subreads were then aligned to respective assemblies with pbmm2 (SMRT Link v12.0.0), a modified version of minimap2,^88^ using parameters “align --preset SUBREAD”. 6mA modifications were identified with “ipdSummary” in kineticsTool (SMRT Link v12.0.0), with parameters “--identify m6A,m4C,m5C_TET --methylFraction”.

To call 6mA bases, we excluded mitochondrial contigs (1 in the MIC, 4 in the MAC assembly) and set a subread coverage threshold of 25 and an identification quality value of ≥ 30 (see Supplementary Methods). Genes ≥1000 bp were selected to assess 6mA levels across gene bodies. The same methods and thresholds were applied to call 6mA in MAC read data of *Blepharisma stoltei*.^12^

### 6mA immunofluorescence

We adapted an existing protocol.^101^ *Loxodes magnus* was harvested, fixed, permeabilized, post-fixed, washed, and resuspended in 3% BSA/TBSTEM as described above. Fixed cells were treated with RNase A (50 µg/mL; 2 h; 37 °C), resuspended in 2 M HCl (20 min; R.T.), washed with 1 M Tris-HCl pH 8, resuspended in 3% BSA/TBSTEM, incubated with primary antibody (Table S5; overnight; 4 °C), washed with BSA/TBSTEM, then incubated with secondary antibody (30 min; R.T.). Cells were counterstained with DAPI, mounted, and imaged as described above.

### Data availability

Software used for this study are available online on GitHub and archived on Zenodo; URLs and DOIs are cited in the main text. Sequencing data are available from the European Nucleotide Archive (ENA): *Loxodes magnus* genomic libraries and assemblies (PRJEB55123), *L. striatus* genomic libraries (PRJEB55752), *L. magnus* nucleosomal DNA libraries (PRJEB55146), *L. magnus* mRNA-seq and sRNA-seq (PRJEB55324). The following data are available from Edmond (Max Planck Digital Library): detailed nuclei purification protocol (https://doi.org/10.17617/3.OYUXDS); flow cytometry run data for nuclei used for genome sequencing (*L. magnus*, https://doi.org/10.17617/3.4THBHC; *L. striatus*, (https://doi.org/10.17617/3.IUFX39), nucleosomal sequencing (*L. magnus*, https://doi.org/10.17617/3.Y18RPV), and Western blotting (*L.magnus*, https://doi.org/10.17617/3.3TQWJX, *L. striatus*, https://doi.org/10.17617/3.GZNWOJ); *L. magnus* genome assemblies and annotations (https://doi.org/10.17617/3.9QTROS); *L. magnus* variant calling and indel annotations (https://doi.org/10.17617/3.NEV8C1); Western blots (https://doi.org/10.17617/3.0DVGMU); immunofluorescence imaging (https://doi.org/10.17617/3.VWAUYE).

## Acknowledgements

We thank Insa Hirschberg and Frank Chan for training and access to the BD FACSMelody; the Max Planck Genome Centre Cologne (https://mpgc.mpipz.mpg.de/home/) for PacBio and RNA-seq library preparation and sequencing; Heike Budde, Christa Lanz, and the Max Planck Institute for Biology Genome Center for additional sequencing; Andre Noll for computer system administration; Abigail Howell and Michael Borg for suggestions to improve the flow sorting protocol; Aurora Panzera, Vanessa Carlos, and Christian Feldhaus for assistance with optical microscopy; Jürgen Berger and Iris Koch for electron microscopy; Sinja Mattes and Amelie Albrecht for culture maintenance; and Klaus Eisler for gift of strains from the former Tübingen teaching collection.

## Author contributions

B.K.B.S.: data curation, formal analysis, investigation, methodology, software, visualization, writing – original draft. A.S.: formal analysis, investigation, methodology, writing – review and editing. D.E.V.: formal analysis, software, writing – review and editing. C.E.: investigation, methodology, writing – review and editing. M.P.: methodology, writing – review and editing. V.S.: methodology, writing – review and editing. B.H.: methodology, resources. E.S.: conceptualization, data curation, formal analysis, funding acquisition, software, supervision, writing – review and editing.

