## Supplementary Information for "Nuclear dualism without extensive DNA elimination in the ciliate *Loxodes magnus*"

### Supplementary Results

*1. Simulation of k-mer comparisons for MIC and MAC of Paramecium tetraurelia*

If the MIC genome contains MIC-limited IESs, which in other ciliates can represent ≥10% of the total MIC sequence content, we would expect to see unique k-mers in the MIC library, with a k-mer frequency peak similar to the main genome peak of shared k-mers. Contamination of the MAC library with MIC sequences would reduce the number of unique k-mers, but the MIC-specific k-mers should still be recognizable as a distinct cluster whose frequencies differ between the MIC and MAC libraries, e.g. in a MIC vs. MAC k-mer frequency heatmap, as long as the target genomes have been enriched.

To explore the expected results of k-mer comparison when libraries have different degrees of contamination (i.e. MAC contaminated with MIC and vice versa), we simulated short-read shotgun sequencing libraries of MIC and MAC genomes from reference assemblies of *Paramecium tetraurelia* strain 51. The *P. tetraurelia* MAC+IES assembly was used in lieu of the de novo MIC assembly because of its higher contiguity. Published real sequencing data from ciliate genomes were not suitable for such benchmarking because the expected purity is unknown, and there is often also contamination from other organisms, e.g. bacteria.

When two pure MAC libraries with high coverage are compared, the only source of unique k-mers should be sequencing errors, visible as a left-sloping curve in the k-mer frequency spectra (Figure S14). Most k-mers in the k-mer comparison heatmap should fall along the

1:1 line (Figure S15). The spectra show multiple peaks because *P. tetraurelia* has experienced several whole genome duplication events in its evolutionary history.

When a pure MAC library is compared against a mixed MAC+MIC library, the k-mers originating from the IESs of the MIC library are noticeable in the frequency curve of k-mers unique to the mixed library (Figure S14, dashed orange lines), which have frequencies above the background sequencing error k-mers. As the fraction of MIC in the mixed library is increased, the frequency peak of these unique k-mers also increases correspondingly. Finally, when a pure MAC library is compared with a pure MIC library, the frequency peaks of both the unique and shared k-mers should align with the overall average k-mer coverage (Figure S14).

In k-mer frequency heatmaps, the MIC-specific k-mers are visible as clusters off the 1:1 axis (Figure S15). Although the difference between MIC and MAC libraries is characterized as MIC-specific IESs, there is also a cluster of MAC-specific k-mers, which corresponds to the sequences at IES junctions after excision.

If both libraries are impure, MIC- and MAC-specific k-mers should nonetheless be detectable as unique k-mers in the k-mer frequency spectra (Figure S16A), and as off-axis clusters in the k-mer comparison heatmaps (Figure S16B), even when only 70% of each library is composed of the target genome.

Despite IESs only comprising 3.6 Mbp total sequence, compared with the total MAC assembly size of 72.1 Mbp (MAC+IES assembly 75.7 Mbp), differences in the two libraries were readily observable in the k-mer comparisons even with moderate cross-contamination.

### *2. rDNA genes are not amplified during development of* Loxodes *MAC genomes*

*Loxodes magnus* assemblies contain multiple loci composed of tandem rDNA arrays. To assess whether rDNA genes may be more amplified in the *L. magnus* MAC genome as it is in ciliates like *Oxytricha trifallax* (56× mean nanochromosome copy number)[1] and

*Tetrahymena thermophila* (200× per MAC genome copy)[2] we mapped all the HiFi CCS reads underlying our genome assemblies to a single, complete, representative rDNA locus (4.9 kbp). rRNA coverage was similar in the MAC (mean 5119, s.d. 385) and MIC (5817, s.d. 347). The total quantity of input CCS reads used in assembly and mapping was 6.6 Gbp and 6.1 Gbp respectively for the MAC and MIC genome libraries. Thus, contrary to the past proposal for *Loxodes*,[3] there is no evidence of amplification of MAC rRNA genes.

#### *3. Putative "IESs" are allelic indel polymorphisms*

Differences between MIC and MAC libraries may not be obvious from high-level k-mer comparison if sorting of nuclei was imperfect, or if IESs constituted only a small percentage of the total genome. Additionally, if the MAC library was contaminated by some MIC sequence, the resulting MAC assembly may include some IESs. However, we reasoned that IESs should still be identifiable from read mappings, so long as there was significant differential enrichment in the samples. They should appear as indels with higher coverage in the MIC than MAC library. Given that the ratio of MIC:MAC in *Loxodes magnus* cells is between 1:1 to 1:2 to begin with, flow sorting should yield significantly >50% of the respective targets in the sorted nuclei libraries. Based on previous analyses of MIC-enrichment libraries of other ciliate species, we expected the following: (a) More IESs should be predicted from the MIC-enriched than the MAC-enriched library. (b) IES retention scores, i.e. IES coverage relative to flanking sequences, should be >0.5 in the MIC library and <0.5 in the MAC library. (c) IESs predicted from the MIC library should be predominantly insertions relative to the MAC reference.

"IESs" from *L. magnus* were probably indel polymorphisms in a diploid genome, because an average retention score of ~0.5 in both MIC and MAC libraries means that reads with vs. without the "IES" have ~1:1 coverage ratio. We first verified that the *L. magnus* genome is diploid heterozygous: single nucleotide polymorphisms (SNPs) called on the MAC assembly were represented by two variants per site at ~1:1 average coverage (Figure S5A, S5B).

"IES" indels in HiFi long reads were often correlated with SNP variants in the same reads, regardless of whether they were from the MIC or MAC libraries (Figure 2C). About 10% of "IES" indels were covered by at least 2 haplotagged reads with and without the indel, and so could be used to test this systematically. Of these, the majority (60-70%) were consistently associated with SNP-based read haplotypes (Figure S5C, S5D), i.e. are likely to represent indel allelic variants. Nonetheless, more indel polymorphisms were bound by terminal direct repeats (TDR) than expected by chance, especially TDRs that contain TA-submotifs (Figure 2E), and hence could have originated from mobile elements.

#### *4. Genome assembly, gene prediction, and genome completeness*

We expected the *Loxodes magnus* genome to have high heterozygosity because the strain used was recently isolated (August 2018) and to our knowledge has not undergone selfing. High repeat content was also expected, from the k-mer spectra of short read libraries. Therefore, we sequenced long-read libraries (PacBio HiFi error-corrected) for de novo assembly of *L. magnus* MIC and MAC genomes.

Long-read assemblies were >10-fold more contiguous than short-read assemblies (N50 ~200 kbp vs. ~10 kbp respectively, Table S1), despite a lower average coverage of 20-30×. Both Flye and Falcon assemblers gave similar contiguity. Because of the expected heterozygosity, we chose the assembly from the diploid-aware Falcon assembler,[4] used a relatively low threshold to collapse heterozygosity, and polished primary contigs with Racon.[5]

201,931 protein-coding genes were predicted in the *L. magnus* MAC reference genome (Figure 3F), excluding predictions that overlapped with low-complexity regions. Of these, 255 contained introns with conflicting orientation and were also excluded. Genes had mean length 949 bp, although the longest prediction of 89 kbp was likely spurious. The majority were single-exon (95%) or had only one intron (3.3%), and most of the empirically defined introns (15,311 of 17,841) were contained within gene predictions. Visual inspection of RNA-seq mapping revealed errors, especially misplaced start or stop sites, which could account for the short average gene length (Figure 3F). Nonetheless, coding sequence appeared to be largely annotated, as the predicted proteome had a completeness of 94% based on 160 of 171 conserved marker genes for Alveolata identified by BUSCO, of which 108 were represented by a single copy. The completeness compared favorably to other published ciliate genomes (83 - 100%, excluding the outlier *Euplotes vannus*; Figure S9).

#### *5. Dicer and Dicer-like proteins in Loxodes magnus*

Dicer proteins (Dcr) are canonically involved in the RNA interference pathway, whereas the related Dicer-like proteins (Dcl) are specific to ciliate genome editing. Both Dcr and Dcl proteins each have a pair of Ribonuclease_III domains (PF00636), but Dcrs have additional N-terminal domains ResIII (PF04851), Helicase_C (PF00271), and Dicer_dimer (PF03368). Like other ciliates, *L. magnus* encodes both Dcrs and Dcls that cluster with their respective counterparts in a tree of the ribonuclease_III domain, and the clusters also correlate with the presence of expected domains (Figure S10).

#### *6. Search for development specific small RNAs in Loxodes magnus*

Ciliate small RNAs (sRNAs) are typically 20-30 nt long,[6–10] and multiple sRNA classes may be expressed, e.g. several classes of small RNAs are produced during *Paramecium* development, two of which, scnRNAs and iesRNAs, are development-specific and involved in IES excision.[7] The shortest class of sRNAs in *Paramecium*, siRNAs (mostly 23 nt), are not development-specific but instead regulate gene expression by silencing.[11,12]

sRNAs from both starved or actively growing *L. magnus* cells had peaks in their length distribution at 24 and 25 nt. A strong 5’ U bias was present in 24 nt sRNAs (81% U), with a weaker one, closer to parity with A in 23, 25 and 26 nt sRNAs (53-55% U; Figure S11A; genomic DNA A=T=37%). This could either be due to overlapping sRNA classes in this size range or to differential, length-dependent processing by the enzymes that produced these sRNAs.

Inspection of contigs with the most mapped 24-26 nt sRNAs revealed clusters predominantly mapping to one strand or the other (e.g. Figure S11B; 24 nt sRNAs — 301 of 304; 25 nt sRNAs — 835 of 839). Thus these sRNAs resemble antisense siRNAs from *Tetrahymena*[13] and *Paramecium*, which are products of an RNA-dependent RNA Polymerase (RdRP),[12] rather than development-specific scnRNAs and iesRNAs which should map to both DNA strands.[7,14] sRNAs in vegetative *L. magnus* cells are likely siRNAs that regulate gene expression rather than assist DNA elimination. However, given the presence of multiple Dicer-like proteins, it is conceivable that *L. magnus* produces conjugation-specific sRNAs with roles unrelated to DNA elimination.

We inspected one cluster of sRNAs with an underlying gene in detail. We selected this cluster because it encodes a protein with convincing homologs detectable by BLASTP in GenBank's nr database of similar length proteins (~620 aa; *L. magnus* gene ID: 000045F.g151_trans; e.g. to GenBank accession: KAJ3017585.1; E-value 7e-118; search conducted on 26.07.2023). The protein this gene encodes has a protein tyrosine and serine/threonine kinase (PFAM:PF07714) domain followed by seven WD-40 repeats (PFAM:PF00400), which presumably would fold into the characteristic propeller structure.[15]

#### *7. Very low expression and paucity of 6mA across retrotransposon sequences*

Despite their abundance, retrotransposon loci in the *Loxodes magnus* MAC had little 6mA (0.04% of ApT vs. 10.9% of ApT in all predicted genes), which is likely due to their extremely low transcription, as observed for *Blepharisma* retrotransposons.[16,17] Genes with reverse transcriptase (PF00078) and endonuclease (PF14529) domains had a median expression of 0 transcripts per million (TPM); mean TPMs were ranked 11th and 13th lowest among 2767 Pfam domains.

#### *8. Misannotation of DNA transposon families by RepeatClassifier*

In comparison to the repeat families annotated as LINEs, those classified as DNA transposons or helitrons (Figure 4B, Table S2) were probably spurious annotations. A putative DNA/Zisupton family did not contain coding sequences with the expected SWIM zinc finger domain (PF04434),[18] whereas putative Unknown/Helitron-2 repeats encoded mostly WD-40 repeat sequences but not the endonuclease or helicase domains characteristic and necessary for helitron replication;[19] RepeatClassifier similarly misclassified abundant WD-40 repeats in *Blepharisma*.[16]

#### *9. Homologs to ISXO2-like transposase in* Loxodes magnus

Two proteins were annotated as "ISXO2-like transposase" (PF12762, DDE_Tnp_IS1595) in *L. magnus* MAC, however these do not appear to be contained in intact mobile elements, unlike in the hypotrich ciliates, where multiple DDE_Tnp_IS1595 transposase genes were found in the MAC genomes of *Oxytricha* [1] and *Stylonychia* [20], and hundreds of insertion sequences (ISs) associated with these transposases were found in the *Oxytricha* MIC genome.[21] Both the MAC- and MIC-encoded *Oxytricha* DDE_Tnp_IS1595 domain-containing genes are also substantially upregulated during development.[1,21] The *L. magnus* DDE_Tnp_IS1595-containing proteins had best BLASTP matches to protein sequences from *Blepharisma* and *Stentor* MAC genomes in GenBank's nr database. This suggested that these proteins may be domesticated transposases acquired before the hetrotrich/karyorelict divergence. *Blepharisma* genes containing the DDE_Tnp_IS1595 domain were upregulated during development like other transposase genes.[17] As judged from a multiple sequence alignment of the *Loxodes* sequences of the seed alignment for PF12762 (using MAFFT v7.450 with E-INS-i algorithm) and inspection of IS1595 family alignments from TnCentral,[22] one of the two sequences appeared to have a complete DDE motif, while the other has DEE. However, there are no signs of IS repeats identified by RepeatMasker flanking these genes in *L. magnus*. Thus it appears that the DDE_Tnp_IS1595 transposase genes in *Loxodes*/*Blepharisma*/*Stentor* may have been acquired independently from those in *Oxytricha* and may no longer be involved in transposition.

#### *10. Cluster of mobile element genes horizontally transferred from* Rickettsia *bacteria to* Loxodes magnus

Few conserved domains related to DNA transposons were detected in the *L. magnus* predicted proteomes. A phage integrase family domain (Pfam PF00589) was found in 28 predicted genes in the MAC genome, but none were associated with interspersed repeats. A YhG-like transposase domain (PF04654) was also found in one predicted protein. This predicted protein encodes an additional domain, "RecT" (PF03837), annotated as DNA-binding, on its N-terminus. BLASTP to GenBank's nr database revealed top hits to both domains, from different *Rickettsia* proteins (E-values of 7e-115 and 1e-81 for PF04654 and PF03837, respectively). The gene encoding this protein (000111F.g160) had sense-strand 6mA characteristic of *L. magnus* DNA (see main text, "*Loxodes magnus* MACs have characteristics of both active chromatin and heterochromatin"), so this gene is probably not from residual contaminating DNA from *Rickettsia* that might have been co-cultured with *L. magnus*. A gene close by (000111F.g16) encodes an integrase (PF13683, "Integrase core domain"). The protein encoded by this gene differs from the phage integrase domain proteins, which have best BLAST matches to *Woesearchaeota*. Additional BLAST searches revealed an ~9.6 kb cluster of six predicted genes (000111F.g159-g164) that all had best matches to *Rickettsia*, so it is likely all were horizontally transferred to *Loxodes* at once.

Since integrases insert DNA, rather than remove and insert it like transposases,[23] it seems unlikely that any of the proteins with integrase domains is involved in editing out pieces of *Loxodes* genomes.

#### *11. Histone H3 homologs in* Loxodes magnus

The *L. magnus* genome encodes multiple histone H3 homologs which cluster into three groups, one among canonical H3 and H3.3 (cluster 1), a divergent group (cluster 2), and one nested among centromeric H3 (cenH3) sequences (cluster 3) (Figure S12). The histone H3 antibody used was raised against an immunogen with 91.7% identity to cluster 1, but only 75% and 50% to clusters 2 and 3 respectively (Abcam Scientific Support, pers. comm.).

Similarly the immunogen for the anti-H3K4me3 antibody had 100% identity to cluster 1 but <65% for clusters 2 and 3, and immunogens for the anti-H3K9ac and H3K9me3 antibodies only matched cluster 1.

#### *12. Homologs of AMT1 methyltransferase complex components in* Loxodes magnus

In predicted proteins from the *L. magnus* MAC genome we detected four clear homologs of the *Tetrahymena* methyltransferase subunits MTA1 (AMT1) and MTA9-B/MTA9 (AMT6/7)[24,25] using BLASTP (E-values 6e-22 to 6e-43). On the other hand, there were no convincing matches (E-value < 1e-3) to *Tetrahymena*'s p1 and p2 proteins (also known as AMTP1 and AMTP2)[26] in *L. magnus* predicted MAC and MIC proteins with BLASTP, whereas there were convincing matches in predicted proteins from the *Blepharisma stoltei* MAC genome (E-value 5e-45 and 4e-16, respectively). We speculate that the p1 and p2 proteins enable complex formation with the methyltransferase subunits, leading to methylation on both strands.

### Supplementary Discussion

*Plausibility of previously reported differential genome amplification and IES excision in* Loxodes

A previous study reported that the copy-number of protein coding genes in an uncultivated *Loxodes* sp. varied across at least four orders of magnitude.[27] This conclusion was based on qPCR quantification of four genes–actin, RS11, EF-1a, and alpha-tubulin–using single-cell multiple-displacement amplification (MDA) products as templates and the SSU rRNA gene as the common reference across samples.

To take the results at face value, we have to assume that: (1) there is little amplification bias from whole genome sequencing, (2) all the genes targeted (except SSU rRNA) are single-copy in the haploid genome, (3) PCR primers are specific despite being degenerate and do not amplify any paralogs or non-target sequences, (4) the four protein-coding genes fortuitously represent the full dynamic range of copy-number variation after differential amplification.

The DNA content of karyorelict MACs is "paradiploid", meaning that it is on the same order of magnitude as the diploid zygotic nucleus precursor.[3,28] Therefore, if there is differential DNA amplification of protein-coding loci in the MAC, there must be a similar amount of DNA that is eliminated during development such that the total amount is about the same. For example, if 10% of the genome is amplified 10-fold, then the remaining 90% must be eliminated.

However, the previous study reported relative abundances spanning 4 orders of magnitude. If we assume that the gene with the lowest abundance is single-copy (i.e. no amplification), that means some loci are amplified $10^4$-fold compared to the zygotic nucleus. To keep the total DNA quantity the same, loci that are amplified $10^4$-fold can constitute a maximum of $10^{-4}$ (0.01%) of the genome, in which case all other DNA must be eliminated.

If we assume that loci with $10^4$-fold amplification are outliers, and that the mean amplification lies between 10- to 100-fold, this still implies that 90% to 99% of the MIC DNA must be eliminated. While on the higher end, this may be consistent with the degree of elimination known from other ciliates, e.g. *Paramecium caudatum* with 1.3 Gbp MIC vs. 30 Mbp MAC genome, i.e. 98% eliminated.[29] Such extensive elimination, however, would be easily detected, even if sort purity was low, by differential k-mer abundances in MIC vs. MAC libraries, which we did not observe.

We therefore judge that differential genome amplification in *Loxodes* to the extent claimed (up to $10^4$-fold) is unlikely. The previous result was based on estimating relative copy number of five genes using qPCR, using multiple-displacement amplification (MDA) products from single cells as the templates. There are multiple paralogs or copies of the genes analyzed by Maurer-Alcala et al. in *Loxodes magnus* as for the rRNA gene, e.g. > 80 copies of actin in both the MAC and MIC genomes. The variability in copy number previously reported is thus a consequence of insufficient knowledge of the underlying genome architecture and gene organization at the time, and may have been compounded by insufficient primer specificity and amplification biases from MDA.

The previous study further concluded that *Loxodes* has IESs by mapping transcriptome data (as a proxy for somatic MAC sequence) to single-cell genome assemblies from MDA products (as a proxy for MIC sequence). They reasoned that because MDA preferentially amplifies longer DNA molecules, it should preferentially amplify MIC DNA because it is less fragmented than MAC, based on a prior study on *Chilodonella uncinata*.[30] They found indels present in the MDA assembly but not the transcriptome; these indels were bound by terminal direct repeats (TDRs), which they took to be a diagnostic feature of IESs in general. Although MDA may preferentially amplify MIC DNA in ciliates with extremely short MAC nanochromosomes like *Chilodonella*, it is doubtful whether this applies for other ciliates that do not have such drastic fragmentation in the MAC; there is no evidence that *Loxodes* or karyorelicts in general have nanochromosomes. In addition to amplification biases mentioned above, MDA is also known to produce chimeric sequence artifacts [31] and hence is not suitable for evaluating genome editing without other supporting evidence. In this study we have observed that allelic indel polymorphisms are often bound by TDRs, so we propose that the putative IESs from the previous study were in fact indel polymorphisms.

### Supplementary Figures

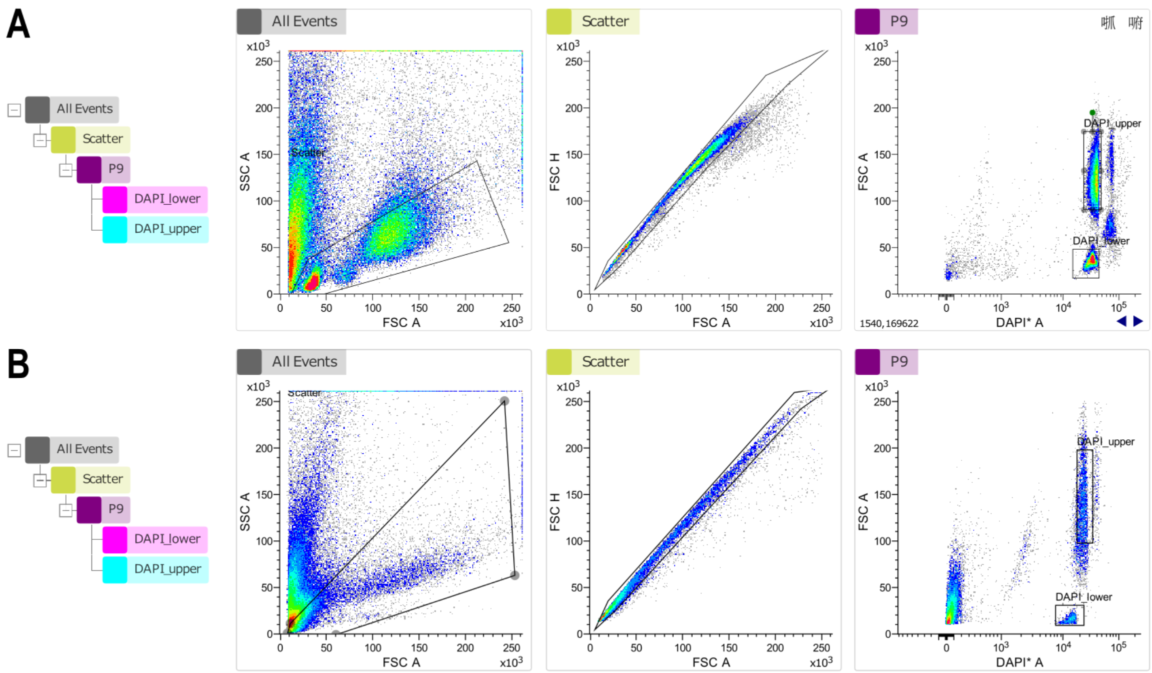


**Figure S1. Gating scheme and scatter plots for fluorescence-activated sorting of *Loxodes* nuclei. (A)** *Loxodes magnus*, **(B)** *Loxodes striatus* (representative runs).

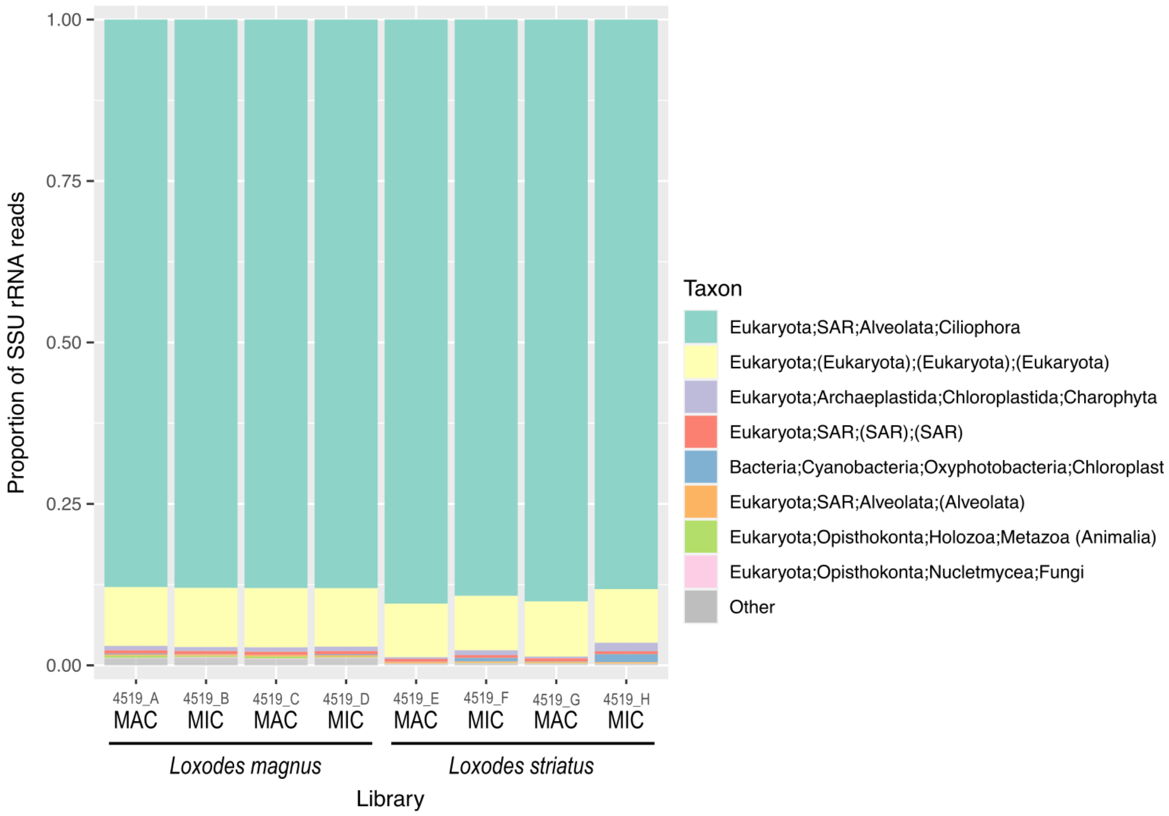


**Figure S2. phyloFlash taxonomic summaries for genomic Illumina libraries of sorted nuclei.** Taxon names in parentheses represent sequences that could not be classified to that rank, the lowest named level was used instead.

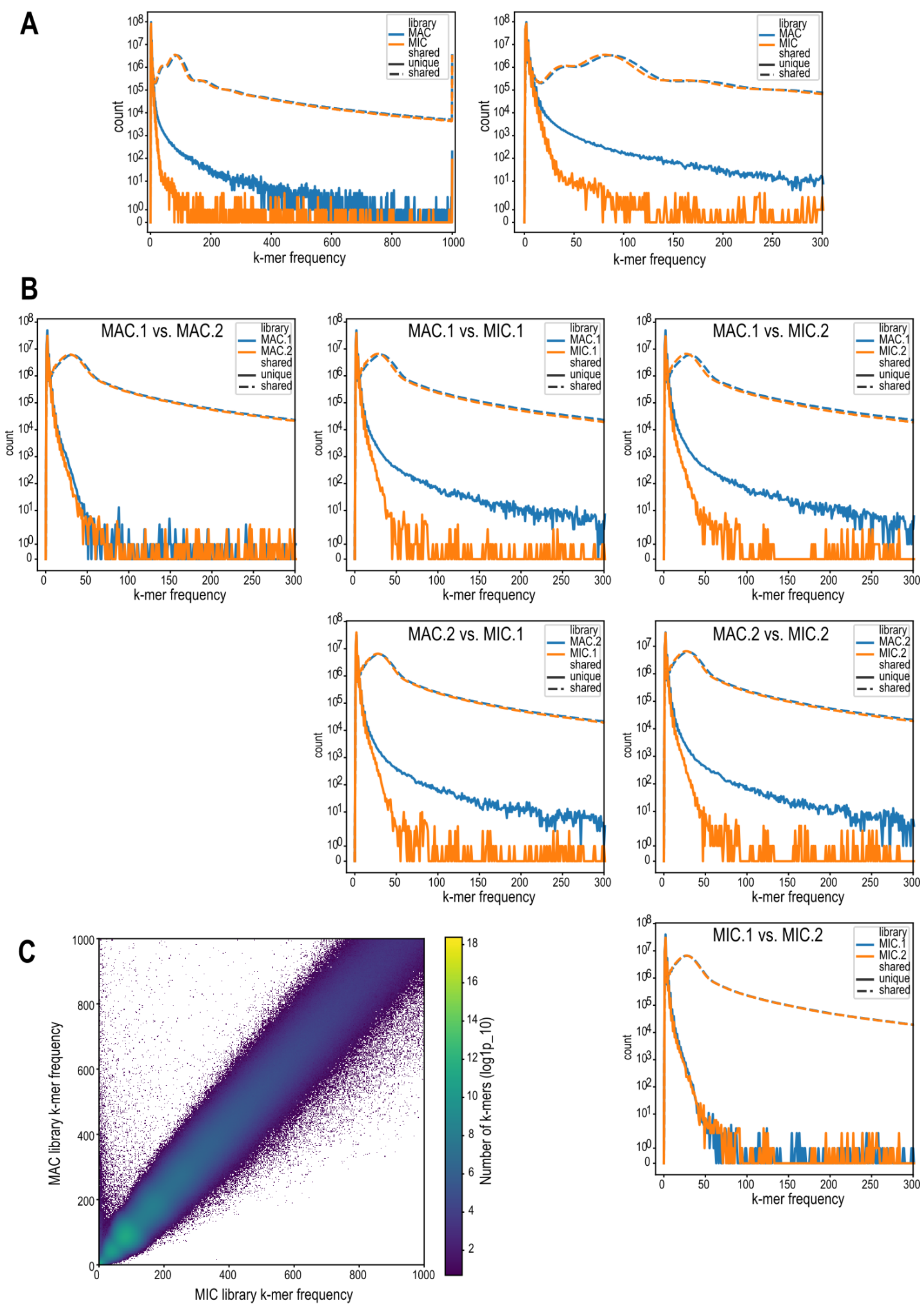


**Figure S3. k-mer comparison of Loxodes magnus MIC vs. MAC libraries. (A)** k-mer multiplicity plot (right: detail) for shared (dashed lines) vs. unique (solid lines) 21-mers in pooled MAC (blue) vs. MIC (orange) sequence libraries. **(B)** k-mer multiplicity plots for shared vs. unique k-mers, pairwise comparison of individual replicates of MAC and MIC, subsamples of 150 M reads per library; x-axis truncated at 300× frequency. **(C)** Heatmap comparing frequency of 21-mers in MIC vs. MAC libraries, axes to 1000× frequency.

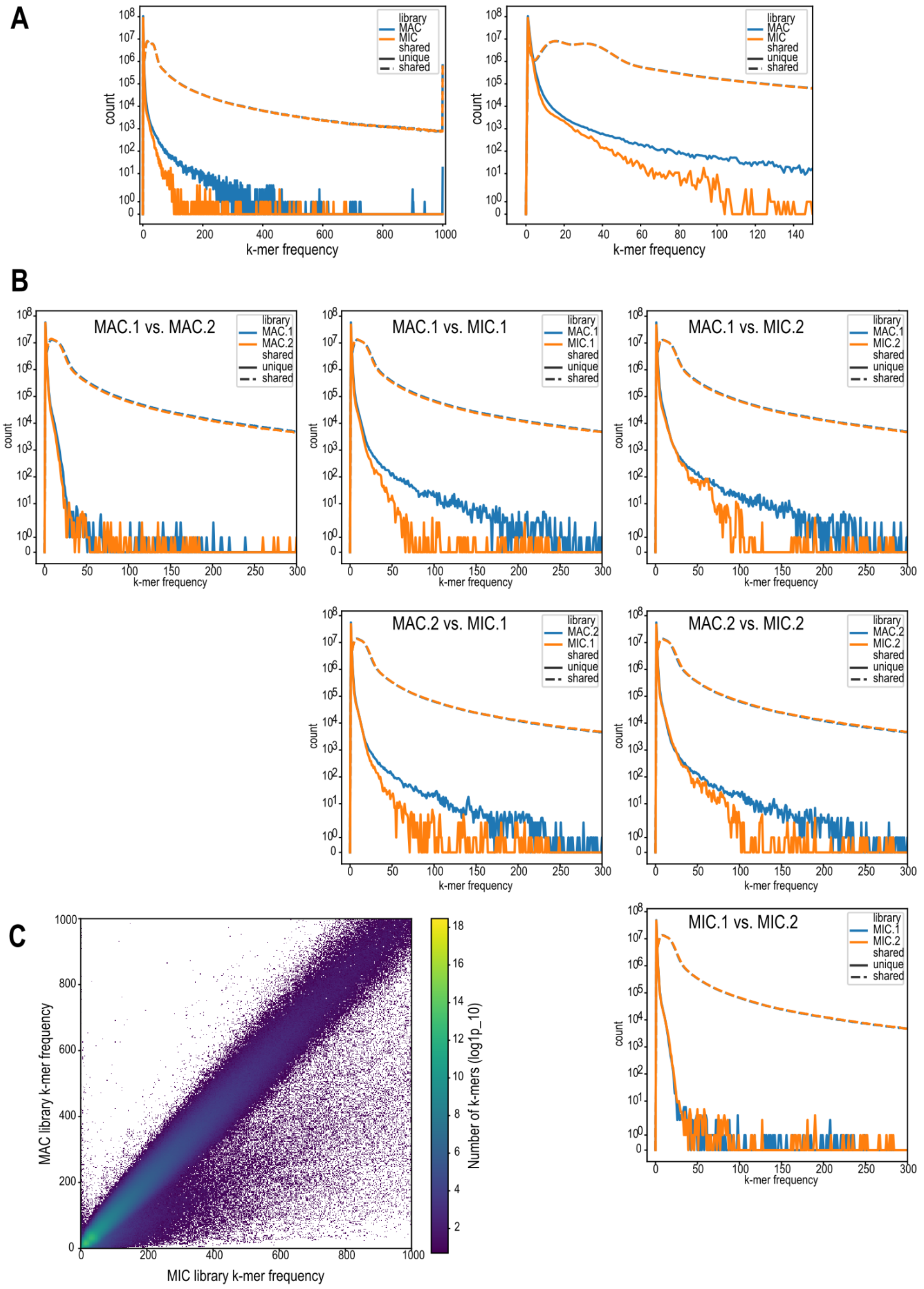


**Figure S4. k-mer comparison of *Loxodes striatus* MIC and MAC genomic libraries.**
Panel captions as for Figure S3.

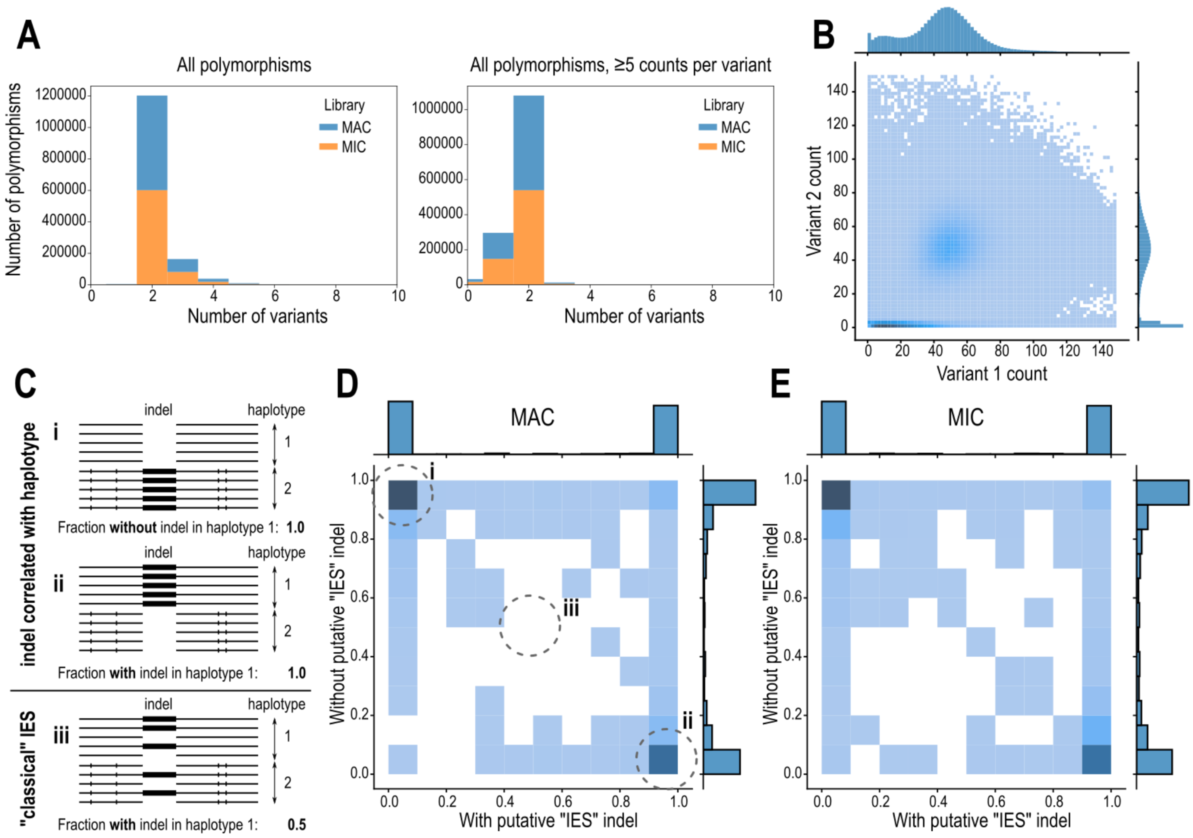


**Figure S5. Single nucleotide polymorphisms (SNPs) in *Loxodes magnus* and correlation with indels. (A)** Number of variants per SNP (naive variant counting), before and after low-abundance variants observed ≤5 times were removed. **(B)** Counts of variant 1 vs. variant 2 per SNP (both MIC and MAC libraries, combined); ratio is ~1:1 on average; SNPs where variant 2 has counts ≤5 are likely to be sequencing or variant-calling errors. **(C)** Diagram of how putative “IES” indels were counted relative to haplotagged reads. Individual HiFi reads were tagged as haplotype 1 or 2 based on SNPs relative to the reference assembly. For each set of reads with or without a given “IES” insert, the numbers of reads per haplotype were counted. **(D, E)** 2-D histograms (linear color intensity scale) of the fraction of reads assigned to reference haplotype for reads with insert (horizontal axis) vs. without insert (vertical axis), for each putative “IES” indel position, for MAC vs. MIC libraries (D, E respectively). In most cases, presence of insert was correlated with the haplotype, as expected if they were allelic indel polymorphisms (panel C scenarios i and ii), whereas classical IESs that have reached fixation would be expected to be independent of haplotype (i.e. in the center of the heatmap, panel C scenario iii).

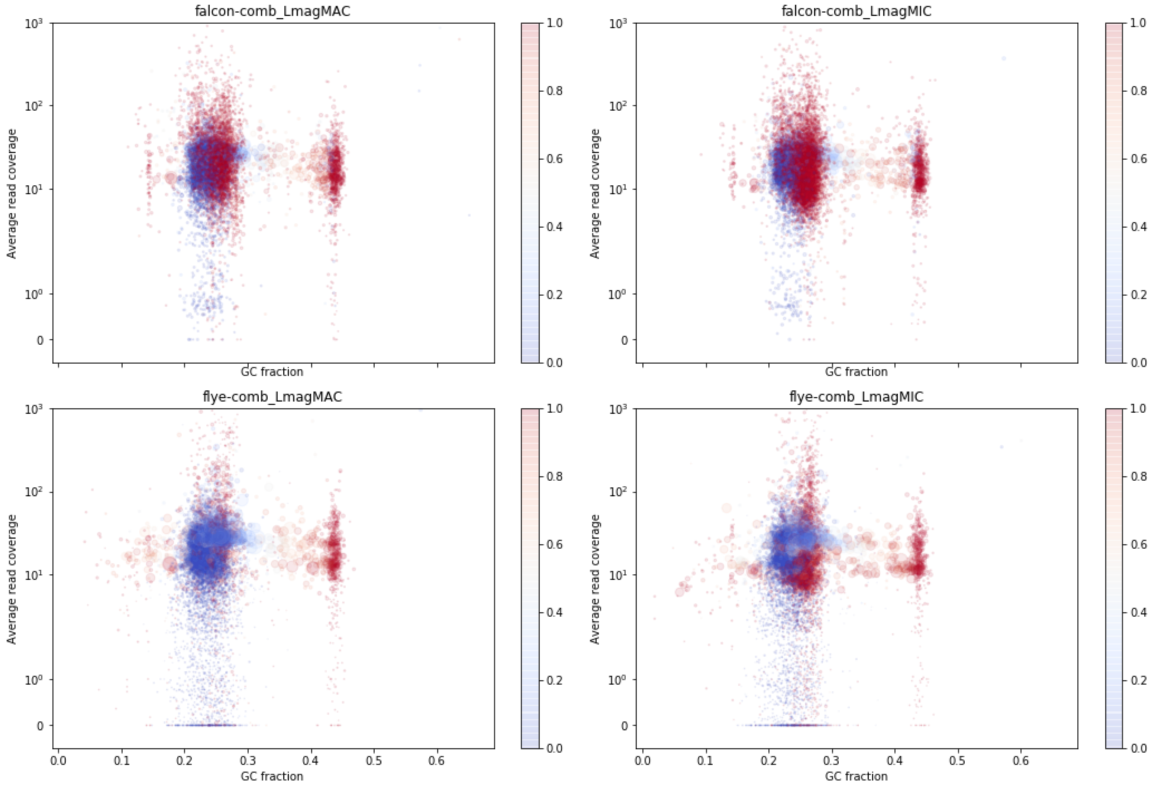


**Figure S6. Blob plots (read coverage vs. GC% per contig) for *Loxodes magnus* long-read genome assemblies.** Plot symbol areas are scaled proportionally to contig length; color scale represents fraction of contig covered by low-complexity tandem repeats.

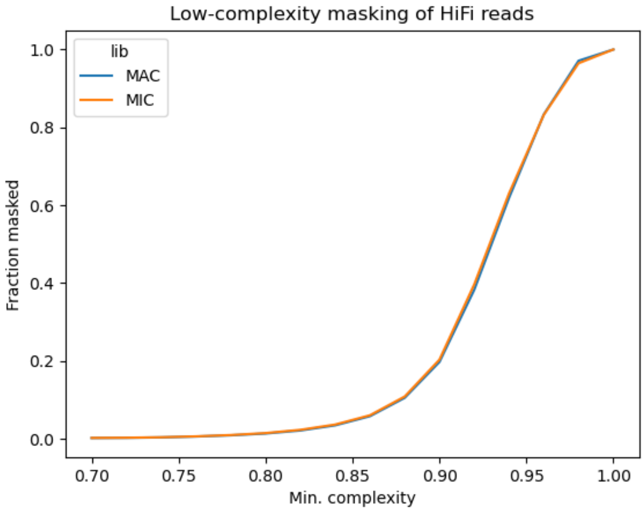


**Figure S7**. **Fraction of bases masked vs. minimum sequence complexity cutoff in masking of low-complexity sequence in *Loxodes magnus* HiFi reads.** Fractions masked at each cutoff level are similar for MICs and MACs, implying that both types of nuclei have similar amounts of low-complexity sequence content.

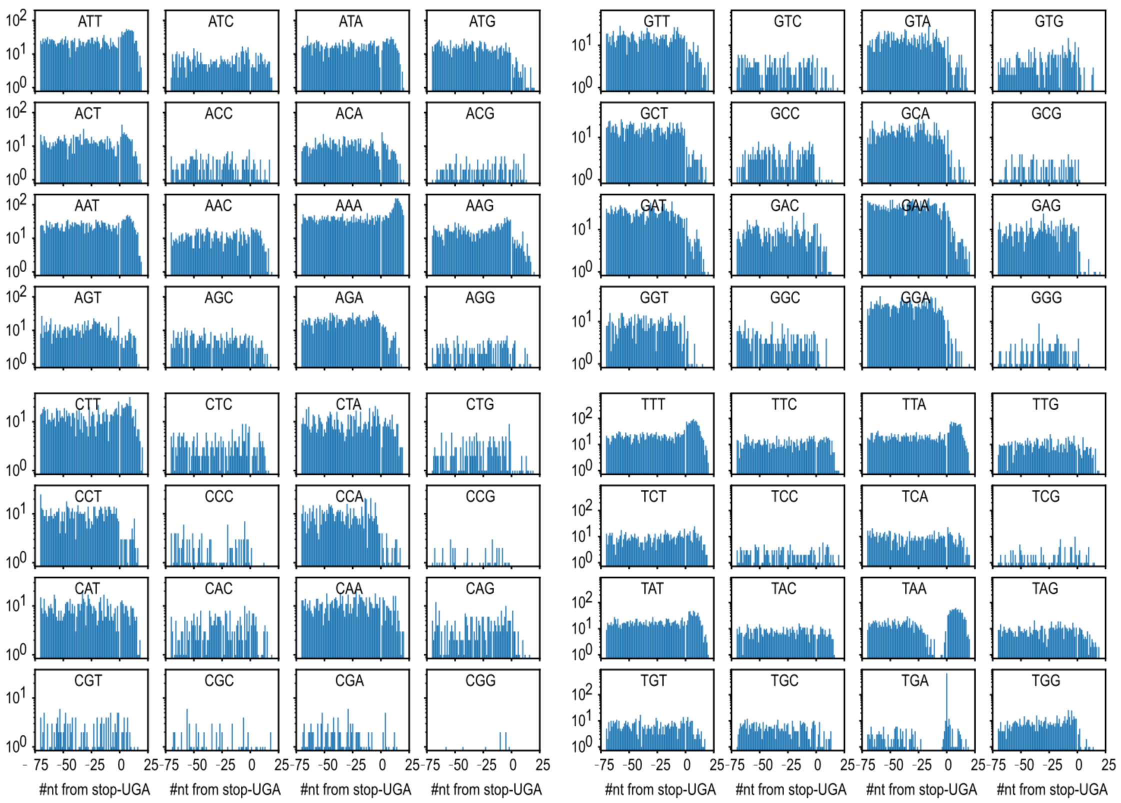


**Figure S8**. **Counts of codons relative to predicted stop-UGAs in *Loxodes magnus* transcripts.**

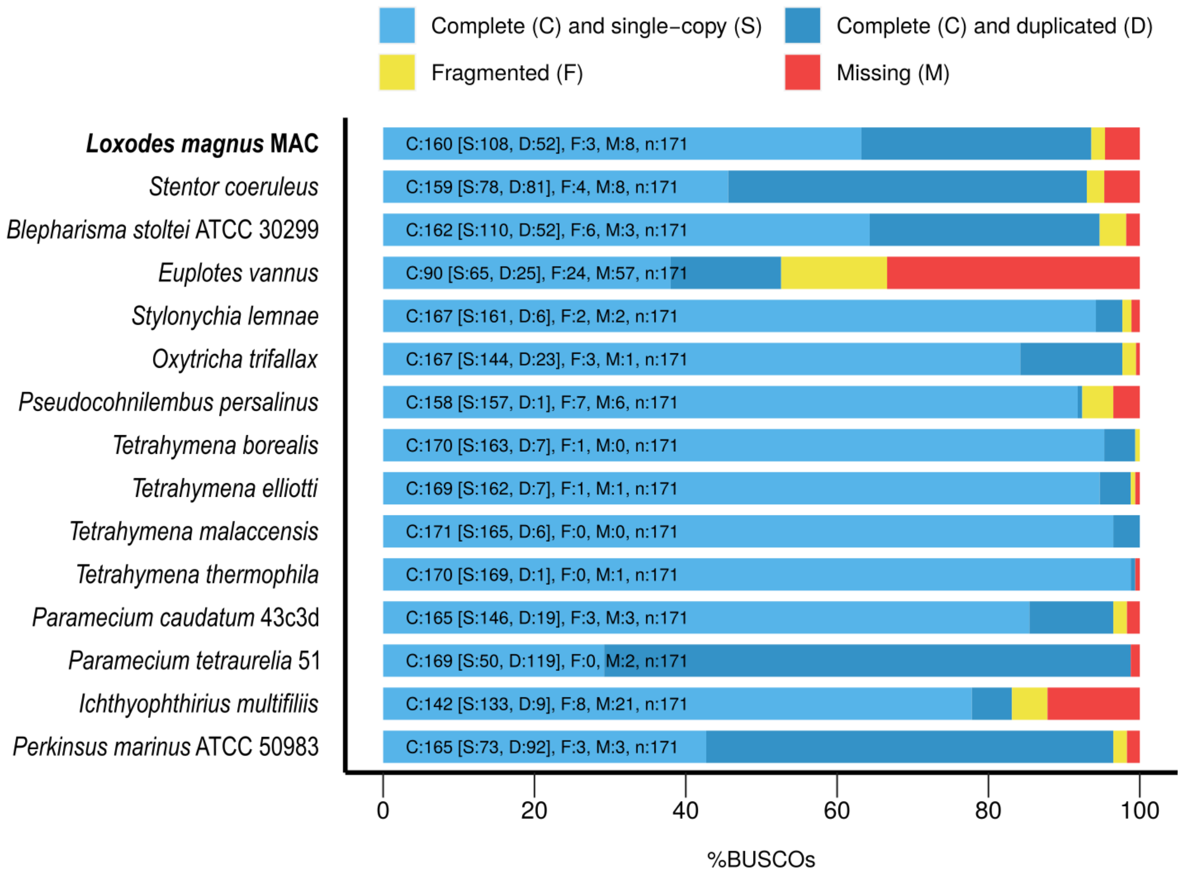


**Figure S9**. **Genome completeness scores using BUSCO conserved orthologs.** Scores calculated using the BUSCO Alveolata marker set, including select ciliates like *Tetrahymena thermophila*. Shown are ciliate MAC genomes (predicted proteins in published annotations) with *Perkinsus marinus* as a relevant non-ciliate, alveolate outgroup.

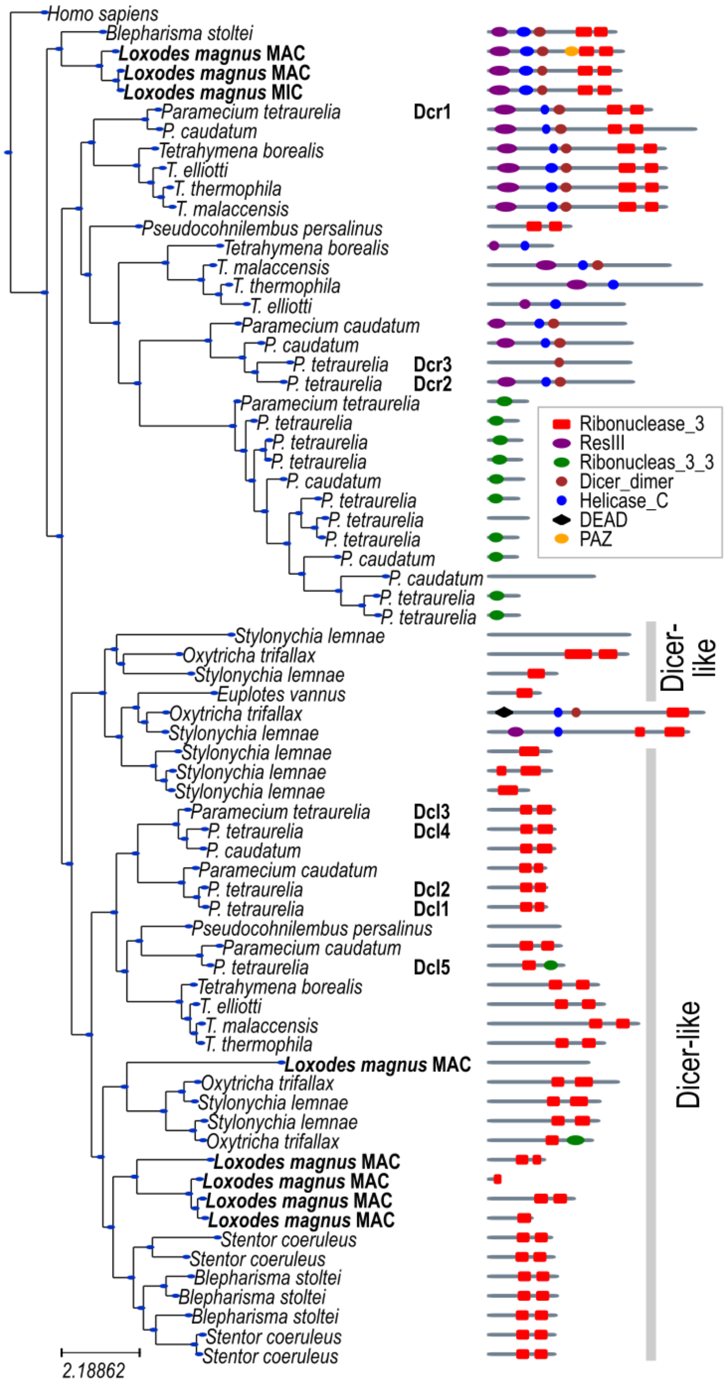


**Figure S10. Phylogenetic tree of Dicer or Dicer-like domains (RNase III domain-like superfamily SSF69065) in ciliate genomes.** Tree shown alongside Pfam domain annotations in the corresponding proteins. Named Dicer (Dcr) or Dicer-like (Dcl) genes in *Paramecium tetraurelia* are indicated.

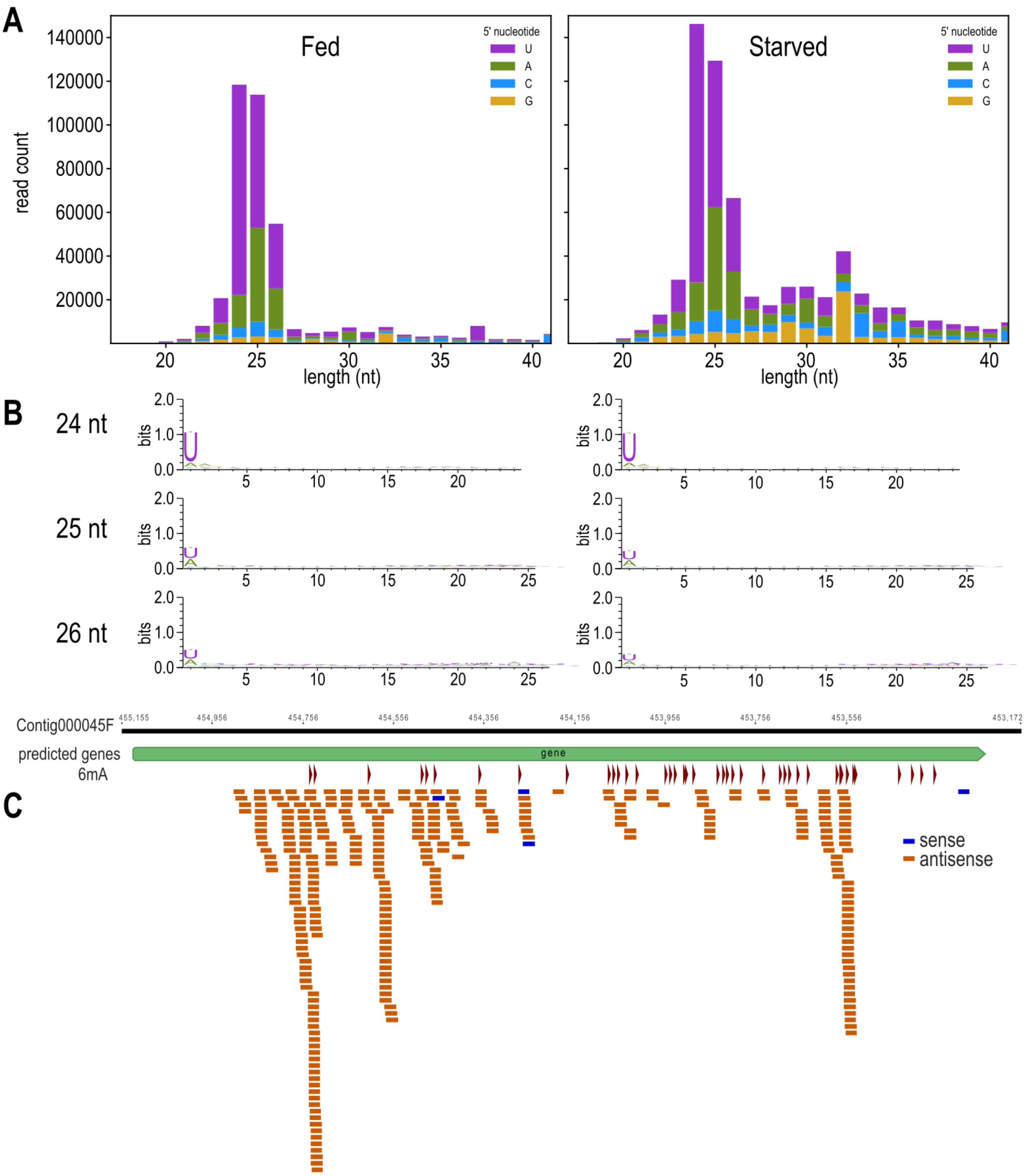


**Figure S11. Characteristics of Loxodes magnus sRNAs. (A)** Length distributions of mapped sRNAs from fed (left) vs. starved (right) populations of *Loxodes magnus*. Bar colors: 5' nucleotide base. **(B)** Sequence logos of 24, 25, and 26 nt sRNAs from fed (left) vs. starved (right) *L. magnus*. **(C)** Mapping of 24 nt sRNAs to a representative *L. magnus* gene, showing strand-bias in both the sRNA (colored bars) and the 6mA modifications (red arrowheads).

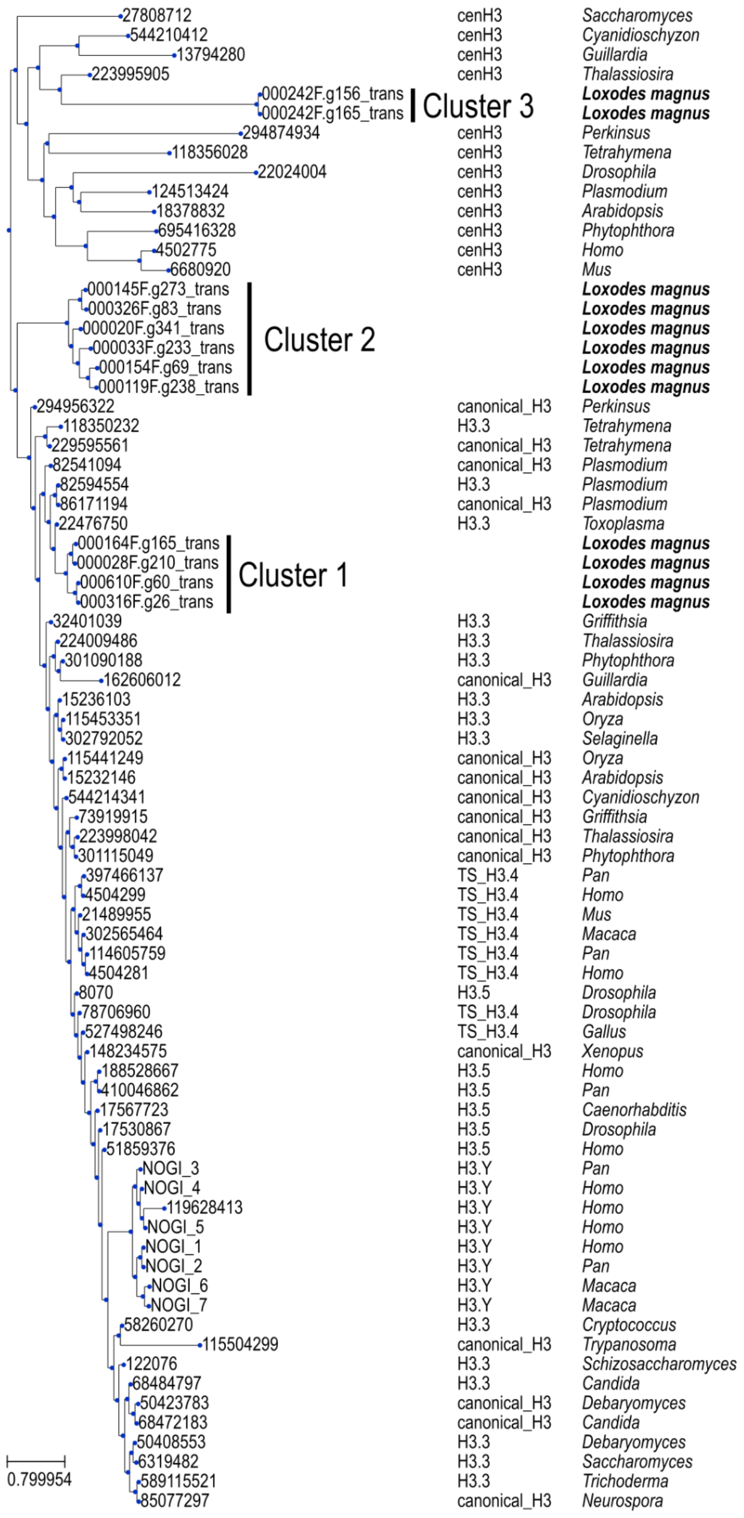


**Figure S12**. **Phylogenetic tree of histone H3 sequences from model organisms and *Loxodes magnus*.** Reference sequences and histone variant classification are from HistoneDB 2.0. Histone H3 sequences from *L. magnus* fall into three clusters (indicated).

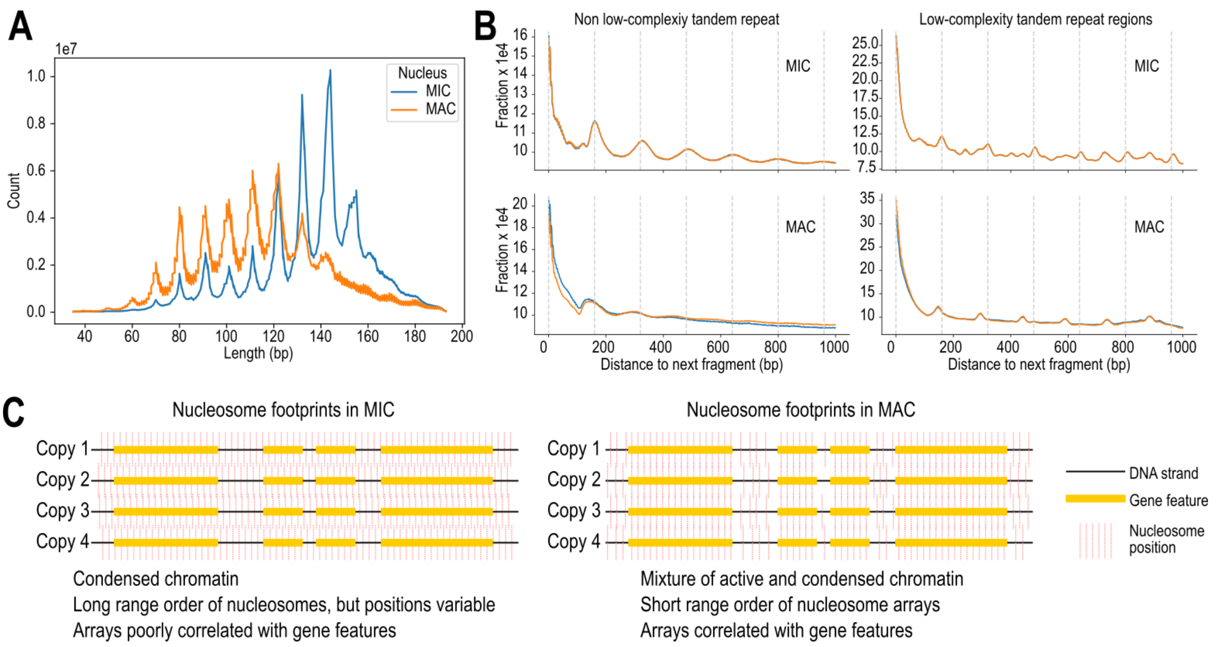


**Figure S13. Properties of *Loxodes* nucleosomes. (A)** Fragment size distribution of dsDNase digest nucleosomal DNA libraries for MIC (blue) vs. MAC (orange), from merged paired-end reads. **(B)** Comparison of global phaseograms of nucleosomal DNA libraries when low complexity repeat regions are masked as in Figure 5D (left), vs. for low complexity repeat regions only (right). **(C)** Schematic model of nucleosome arrays in Loxodes MIC vs. MAC nuclei.

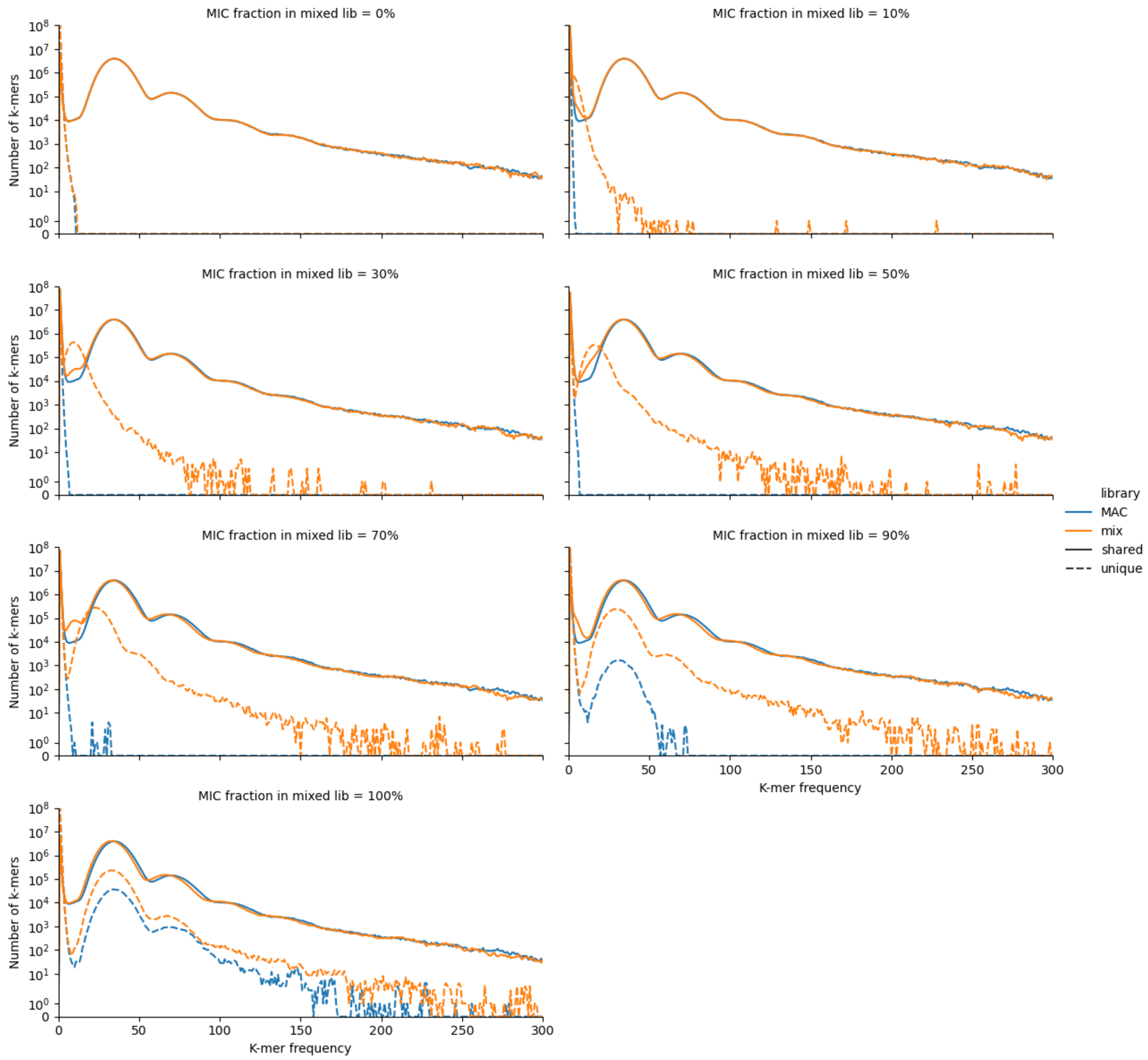


**Figure S14. k-mer comparisons (k=19) of simulated pure MAC library vs. MAC+MIC mixture.** Simulations with increasing percentage of MIC ("mix"). Solid lines – shared k-mers, dashed lines – unique k-mers, blue – pure MAC library, orange – mixed library.

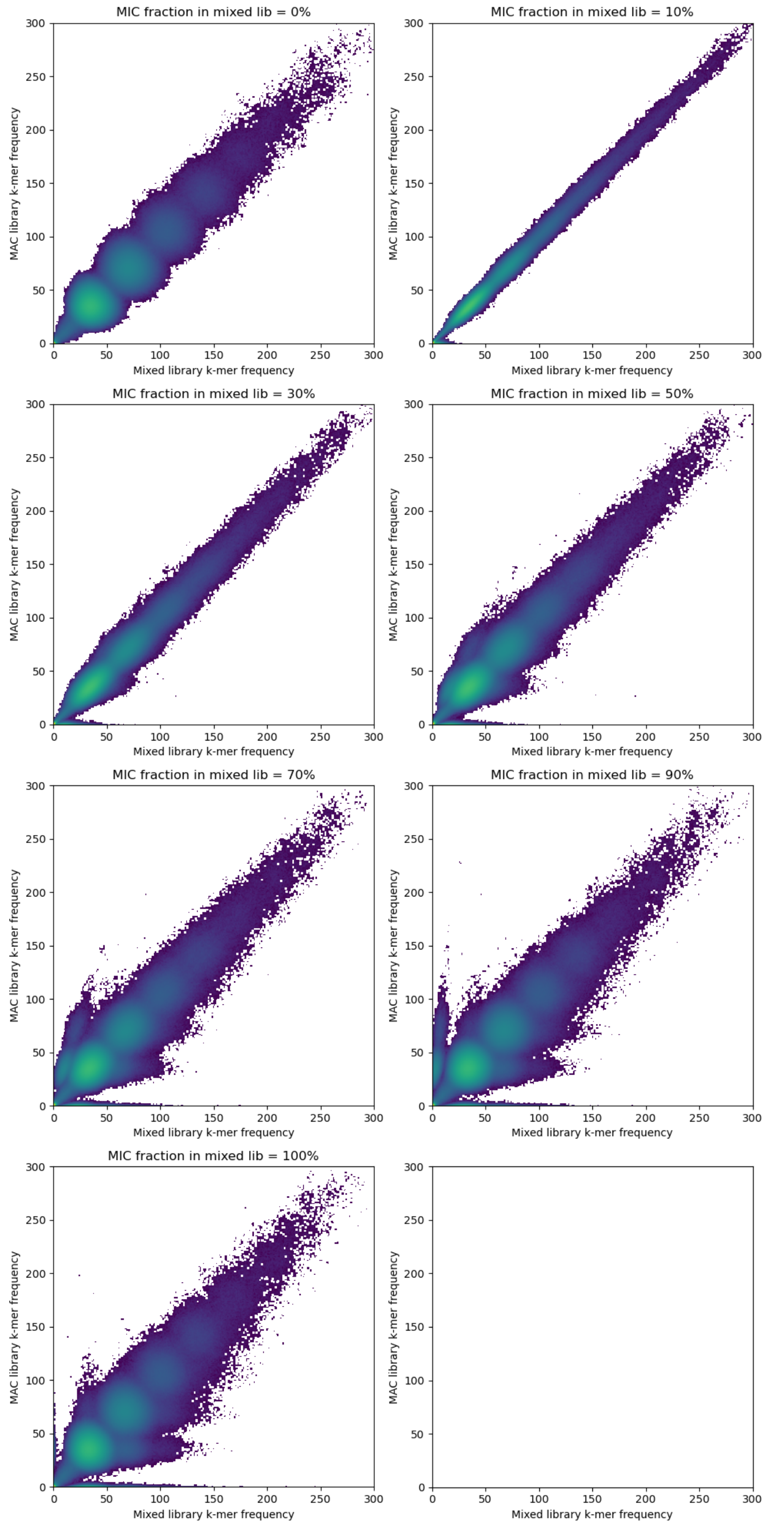


**Figure S15. Heatmaps of k-mer frequency in comparisons of simulated pure MAC library (vertical axis) vs. MAC+MIC mixture.** Ordered by increasing percentage of MIC (horizontal axis); color scale intensity represents number of k-mers.

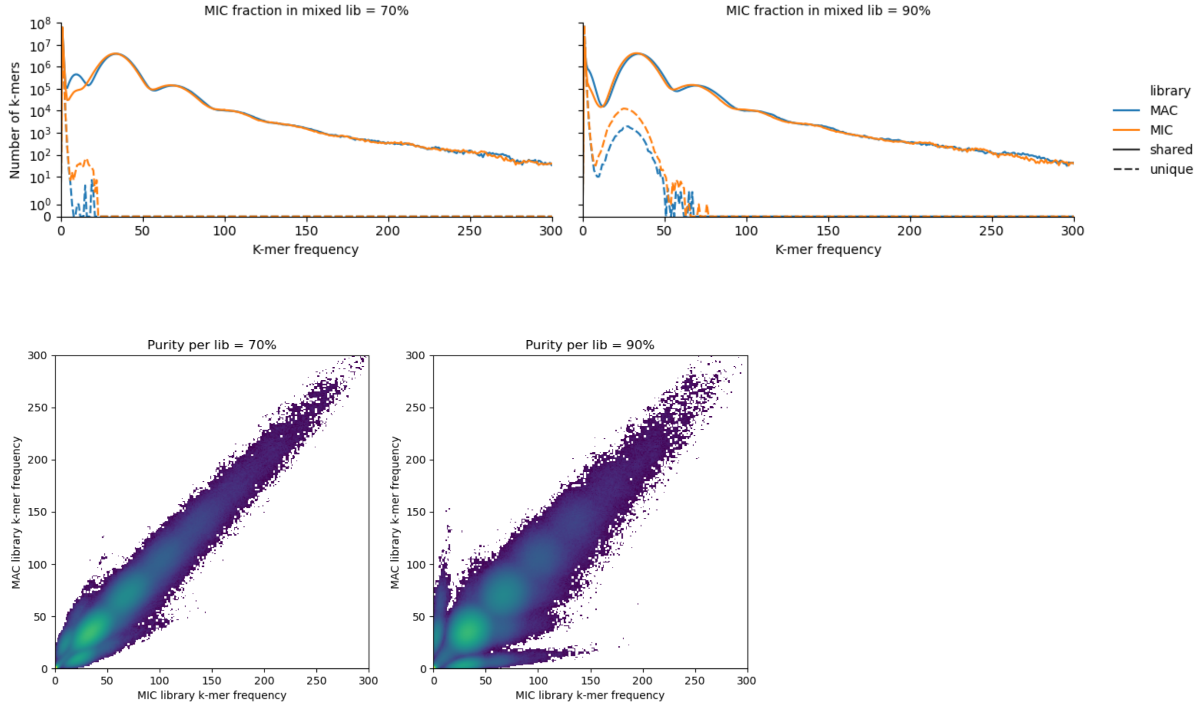


**Figure S16. Simulated k-mer contamination. (A)** K-mer comparisons (k=19) of simulated MAC vs. MIC libraries, each with cross-contamination from the other sequence type. Left: both libraries 70% pure, right: both libraries 90% pure. **(B)** Heatmaps of k-mer frequency in comparisons of simulated cross-contaminated MAC vs. MIC libraries.

### Supplementary Tables

**Table S1.** Genome assembly metrics for different assemblers and sorted nuclei libraries. Low complexity regions were merged if overlapping, and only regions >1 kbp were counted. Interspersed repeat total length: after overlapping annotations were merged. ND – not determined.

| **Assembler** | **Falcon** | **Flye** | **MEGAhit** | **SPAdes** | **Falcon** | **Flye** | **MEGAhit** | **SPAdes** |
|---|---|---|---|---|---|---|---|---|
| **Nucleus** | **MAC** | **MAC** | **MAC** | **MAC** | **MIC** | **MIC** | **MIC** | **MIC** |
| **Library type** | **PacBio HiFi** | PacBio HiFi | Illumina 2×150 bp | Illumina 2×150 bp | **PacBio HiFi** | PacBio HiFi | Illumina 2×150 bp | Illumina 2×150 bp |
| **Number contigs** | **7858** | 8222 | 619582 | 3287736 | **9387** | 9231 | 602227 | 3459997 |
| **Total length (Mbp)** | **706** | 626 | 404 | 793 | **848** | 805 | 399 | 815 |
| **Largest contig (Mbp)** | **2.89** | 2.77 | 0.392 | 0.450 | **2.867** | 2.98 | 0.509 | 0.502 |
| **GC (%)** | **26.2** | 25.91 | 24.65 | 24.45 | **26.1** | 25.89 | 24.65 | 24.48 |
| **N50 (kbp)** | **177** | 204 | 10.2 | 6.59 | **193** | 244 | 10.3 | 6.57 |
| **N75 (kbp)** | **65.4** | 75.8 | 2.34 | 1.61 | **68.6** | 93.4 | 2.33 | 1.60 |
| **L50** | **802** | 647 | 3772 | 6997 | **922** | 749 | 3706 | 6983 |
| **L75** | **2476** | 1954 | 17488 | 30669 | **2809** | 2116 | 17420 | 30757 |
| **# N's per 100 kbp** | **0** | 3.43 | 0 | 70.94 | **0** | 3.68 | 0 | 63.4 |
| **Low complexity (Mbp)** | **231** | 172 | ND | ND | **359** | 344 | ND | ND |
| **Interspersed repeats (Mbp)** | **454** | ND | ND | ND | **571** | ND | ND | ND |
| **Non-repetitive (Mbp)** | **229** | ND | ND | ND | **245** | ND | ND | ND |

**Table S2**. Summary of RepeatMasker annotations in MAC vs. MIC genome assemblies for interspersed repeat families found by RepeatModeler with a classification to known mobile element class, out of a total of 170 putative repeat families. Full length copies are defined as annotations >80% and <120% of consensus length.

| Repeat family | Classification | Cons. length (bp) | Total annotated length (bp) | | No. copies | | No. full length copies | |
|---|---|---|---|---|---|---|---|---|
| | | | MAC | MIC | MAC | MIC | MAC | MIC |
| rnd-1_family-2 | Unknown/Helitron-2 | 3032 | 64725198 | 61022801 | 2777 | 2805 | 164 | 163 |
| rnd-6_family-790 | Unknown/Helitron-2 | 13725 | 5659472 | 5410868 | 2623 | 2564 | 57 | 63 |
| rnd-1_family-27 | LINE/RTE-X | 6042 | 4861662 | 4818431 | 2816 | 2870 | 311 | 294 |
| rnd-1_family-19 | LINE/RTE-BovB | 5380 | 4183278 | 4405536 | 3132 | 3317 | 313 | 339 |
| rnd-1_family-63 | LINE | 5979 | 3393684 | 3384063 | 3325 | 3402 | 138 | 135 |
| rnd-1_family-12 | LINE/RTE-X | 5327 | 3236685 | 3462892 | 3644 | 3928 | 74 | 74 |
| rnd-1_family-8 | Unknown/Helitron-2 | 6643 | 2946027 | 2767078 | 1412 | 1375 | 83 | 72 |
| rnd-1_family-60 | LINE/RTE-X | 5950 | 2784869 | 2675233 | 1765 | 1777 | 159 | 151 |
| rnd-1_family-18 | LINE/RTE-BovB | 3770 | 2440970 | 2536625 | 5610 | 5844 | 77 | 84 |
| rnd-1_family-5 | Unknown/Helitron-2 | 963 | 2315210 | 2136144 | 665 | 696 | 214 | 218 |
| rnd-1_family-80 | LINE/RTE-X | 3159 | 2099005 | 2276340 | 1804 | 1924 | 318 | 335 |
| rnd-4_family-203 | Unknown/Helitron-2 | 7805 | 2052133 | 2058648 | 1794 | 1829 | 48 | 46 |
| rnd-1_family-35 | LINE/RTE-X | 3420 | 1986760 | 2079082 | 1720 | 1825 | 224 | 237 |
| rnd-1_family-87 | LINE/RTE-BovB | 4907 | 1519913 | 1749801 | 2380 | 2588 | 29 | 32 |
| rnd-5_family-51 | LINE/CR1-Zenon | 3464 | 1082459 | 1154752 | 887 | 954 | 123 | 129 |
| rnd-1_family-284 | LINE | 2171 | 913602 | 907091 | 1010 | 1007 | 227 | 217 |
| rnd-1_family-9 | Unknown/Helitron-2 | 2032 | 804242 | 867379 | 2753 | 2925 | 64 | 76 |
| rnd-1_family-3 | Unknown/Helitron-2 | 1055 | 767788 | 713631 | 791 | 797 | 400 | 394 |
| rnd-5_family-124 | LINE/Proto2 | 1101 | 658646 | 670670 | 1160 | 1186 | 350 | 357 |
| rnd-1_family-409 | LINE/CR1-Zenon | 2825 | 657715 | 655773 | 460 | 483 | 133 | 116 |
| rnd-1_family-98 | Unknown/Helitron-2 | 1233 | 646506 | 704277 | 469 | 503 | 23 | 30 |
| rnd-1_family-42 | Unknown/Helitron-2 | 1768 | 571872 | 538255 | 425 | 432 | 48 | 44 |
| rnd-1_family-395 | LINE | 1041 | 533381 | 569970 | 637 | 668 | 465 | 499 |
| rnd-1_family-69 | LINE/RTE-X | 2717 | 500464 | 473997 | 373 | 372 | 121 | 118 |
| rnd-6_family-684 | Unknown/Helitron-2 | 4143 | 382572 | 364527 | 561 | 562 | 19 | 15 |
| rnd-4_family-192 | LINE/RTE-BovB | 5540 | 285209 | 302764 | 1394 | 1473 | 10 | 9 |
| rnd-6_family-221 | LTR/Pao | 5722 | 261010 | 381599 | 353 | 532 | 4 | 4 |
| rnd-1_family-61 | LINE | 2131 | 257315 | 272059 | 507 | 517 | 2 | 1 |
| rnd-1_family-112 | LINE/RTE-X | 2037 | 176947 | 183977 | 378 | 395 | 3 | 3 |
| rnd-5_family-642 | DNA/Zisupton | 1851 | 161878 | 162381 | 164 | 165 | 34 | 37 |
| rnd-1_family-125 | LTR/Copia | 281 | 152222 | 156034 | 655 | 675 | 431 | 441 |
| rnd-1_family-154 | LINE | 1997 | 146969 | 134998 | 253 | 243 | 13 | 10 |

| | | | | | | | | |
|---|---|---|---|---|---|---|---|---|
| rnd-1_family-10 | Unknown/Helitron-2 | 320 | 116490 | 67605 | 191 | 180 | 13 | 14 |
| rnd-1_family-212 | Unknown/Helitron-2 | 1188 | 66226 | 65841 | 132 | 142 | 8 | 6 |
| rnd-1_family-94 | Unknown/Helitron-2 | 546 | 38712 | 41004 | 131 | 137 | 10 | 8 |
| rnd-1_family-133 | LINE/RTE-X | 888 | 22562 | 26696 | 56 | 61 | 1 | 2 |
| rnd-1_family-436 | LINE/RTE-RTE | 445 | 20925 | 19592 | 65 | 67 | 35 | 30 |
| rnd-1_family-123 | LINE/R2 | 316 | 12516 | 14397 | 96 | 111 | 9 | 10 |
| rnd-1_family-384 | LTR/Pao | 146 | 2889 | 2758 | 22 | 21 | 7 | 7 |
| rnd-1_family-40 | Unknown/Helitron-2 | 468 | 1395 | 1741 | 8 | 7 | 1 | 2 |
| rnd-1_family-142 | LINE | 317 | 1306 | 1191 | 9 | 9 | 1 | 1 |

**Table S3**. Cell counts of starved vs. fed cultures used for RNAseq of non-dividing vs. actively dividing populations. Three counts (0.1 mL aliquots each) were taken per replicate per time point. Each sample had a total volume of 150 mL. Fed cultures were fed on days 4 and 5.

| Sample | Condition | Date | Time | Count 1 | Count 2 | Count 3 | Density (cells/mL) |
|---|---|---|---|---|---|---|---|
| Lm5-FedA | fed | 2019-08-27 | 16:15:00 | 31 | 27 | 21 | 263.3 |
| Lm5-FedB | fed | 2019-08-27 | 16:15:00 | 26 | 29 | 19 | 246.7 |
| Lm5-FedC | fed | 2019-08-27 | 16:15:00 | 28 | 30 | 19 | 256.7 |
| Lm5-FedA | fed | 2019-08-28 | 14:15:00 | 22 | 23 | 23 | 226.7 |
| Lm5-FedB | fed | 2019-08-28 | 14:15:00 | 23 | 28 | 21 | 240 |
| Lm5-FedC | fed | 2019-08-28 | 14:15:00 | 27 | 25 | 20 | 240 |
| Lm5-FedA | fed | 2019-08-29 | 11:30:00 | 17 | 25 | 19 | 203.3 |
| Lm5-FedB | fed | 2019-08-29 | 11:30:00 | 20 | 21 | 23 | 213.3 |
| Lm5-FedC | fed | 2019-08-29 | 11:30:00 | 20 | 19 | 21 | 200 |
| Lm5-FedA | fed | 2019-08-30 | 10:50:00 | 23 | 22 | 18 | 210 |
| Lm5-FedB | fed | 2019-08-30 | 10:50:00 | 22 | 21 | 16 | 196.7 |
| Lm5-FedC | fed | 2019-08-30 | 10:50:00 | 29 | 34 | 30 | 310 |
| Lm5-FedA | fed | 2019-08-31 | 12:00:00 | 19 | 31 | 22 | 240 |
| Lm5-FedB | fed | 2019-08-31 | 12:00:00 | 29 | 24 | 32 | 283.3 |
| Lm5-FedC | fed | 2019-08-31 | 12:00:00 | 30 | 45 | 26 | 336.7 |
| Lm5-StrA | starved | 2019-08-27 | 16:15:00 | 29 | 21 | 24 | 246.7 |
| Lm5-StrB | starved | 2019-08-27 | 16:15:00 | 27 | 36 | 23 | 286.7 |
| Lm5-StrC | starved | 2019-08-27 | 16:15:00 | 24 | 21 | 26 | 236.7 |
| Lm5-StrA | starved | 2019-08-28 | 14:15:00 | 22 | 25 | 16 | 210 |
| Lm5-StrB | starved | 2019-08-28 | 14:15:00 | 14 | 24 | 33 | 236.7 |
| Lm5-StrC | starved | 2019-08-28 | 14:15:00 | 26 | 28 | 24 | 260 |
| Lm5-StrA | starved | 2019-08-29 | 11:30:00 | 20 | 27 | 19 | 220 |
| Lm5-StrB | starved | 2019-08-29 | 11:30:00 | 18 | 23 | 30 | 236.7 |
| Lm5-StrC | starved | 2019-08-29 | 11:30:00 | 25 | 24 | 31 | 266.7 |

**Table S4**. Published ciliate predicted proteomes (for MAC) or genome assemblies (for MIC) used for comparisons of genome completeness and functional annotations.

| Species | Version | URL | Publication |
|---|---|---|---|
| *Blepharisma stoltei* ATCC 30299 MAC | | https://bleph.ciliate.org/common/downloads/bleph/Bsto_ATCC_protein.fasta | [17] |
| *Blepharisma stoltei* ATCC 30299 MAC+IES | | https://bleph.ciliate.org/common/downloads/bleph/bsto_atcc_mac_plus_ies.fa | [16] |
| *Euplotes vannus* MAC | Mar 2018 | https://evan.ciliate.org/common/downloads/evan/Euplotes_vannus_Mar2018_proteins.fasta | [32] |
| *Ichthyophthirius multifiliis* MAC | | https://ich.ciliate.org/common/downloads/ich/img1_0407.aa.fsa | [33] |
| *Oxytricha trifallax* MAC | 022112 | https://oxy.ciliate.org/common/downloads/oxy/Oxytricha_trifallax_022112_aa.fasta | [1] |
| *Oxytricha trifallax* MIC | | https://oxy.ciliate.org/common/downloads/oxy/Oxytricha_trifallax_micronuclear_assembly.fasta | [21] |
| *Paramecium caudatum* 43c3d MAC | v2.0 | https://paramecium.i2bc.paris-saclay.fr/files/Paramecium/caudatum/43c3d/annotations/caudatum_43c3d_assembly_v1/pcaudatum_43c3d_annotation_v2.0.protein.fa | [34] |
| *Paramecium tetraurelia* strain 51 MAC | v2.0 | https://paramecium.i2bc.paris-saclay.fr/files/Paramecium/tetraurelia/51/annotations/ptetraurelia_mac_51/ptetraurelia_mac_51_annotation_v2.0.protein.fa | [34] |
| *Paramecium tetraurelia* strain 51 MAC+IES | v1.0 | https://paramecium.i2bc.paris-saclay.fr/files/Paramecium/tetraurelia/51/sequences/ptetraurelia_mac_51_with_ies.fa | [34] |
| *Perkinsus marinus* ATCC 50983 (outgroup) | GCF_000006405 | https://ftp.ncbi.nlm.nih.gov/genomes/all/GCF/000/006/405/GCF_000006405.1_JCVI_PMG_1.0/GCF_000006405.1_JCVI_PMG_1.0_protein.faa.gz | |
| *Pseudocohnilembus persalinus* MAC | GCA_001447515 | https://ftp.ncbi.nlm.nih.gov/genomes/all/GCA/001/447/515/GCA_001447515.1_ASM144751v1/GCA_001447515.1_ASM144751v1_protein.faa.gz | [35] |
| *Stentor coeruleus* MAC | Nov 2017 | https://stentor.ciliate.org/common/downloads/stentor/S_coeruleus_Nov2017_proteins.fasta | [36] |
| *Stylonychia lemnae* MAC | Nov 2017 | https://stylo.ciliate.org/common/downloads/stylo/stylo_protein.fa | [20] |
| *Tetrahymena borealis* MAC | Oct 2012 | https://tet.ciliate.org/common/downloads/tet/T_borealis_oct2012_proteins.fasta | [37] |
| *Tetrahymena elliotii* MAC | Oct 2012 | https://tet.ciliate.org/common/downloads/tet/T_elliotti_oct2012_proteins.fasta | [37] |

| | | | |
|---|---|---|---|
| *Tetrahymena malaccensis* MAC | Oct 2012 | https://tet.ciliate.org/common/downloads/tet/T_malaccensis_oct2012_proteins.fasta | [37] |
| *Tetrahymena thermophila* MAC | Mar 2020 | https://tet.ciliate.org/common/downloads/tet/T_thermophila_mar2020-Protein%20fasta.fasta | [38] |
| *Tetrahyemna thermophila* MIC | 2016 | https://tet.ciliate.org/common/downloads/tet/2016_mic.genome.fasta | [39] |

**Table S5**. Primary antibodies used in this study.

| Target | Supplier | Catalog no. | Type | IF dilution (1º) | IF dilution (2º) | WB dilution (1º) | WB dilution (2º) |
|---|---|---|---|---|---|---|---|
| H3K9ac | Merck | 06-942-S | rabbit polyclonal IgG | 1:100 | 1:200 | 1:3000 | 1:5000 |
| H3K9me3 | Merck | 07-442 | rabbit polyclonal IgG | 1:100 | 1:2000 | 1:3000 | 1:5000 |
| H3K4me3 | Abcam | ab8580 | rabbit polyclonal IgG | 1:100 | 1:200 | 1:5000 | 1:5000 |
| Histone H3 | Abcam | ab1791 | rabbit polyclonal IgG | 1:100 | 1:200 | 1:3000 | 1:5000 |
| Histone H4 | Abcam | ab177840 | rabbit monoclonal IgG | 1:100 | 1:200 | 1:500 | 1:2500 |
| 6mA | Synaptic Systems | 202 003 | rabbit polyclonal IgG | 1:2000 | 1:200 | - | - |

### Supplementary Methods

#### *Control against inadvertent cross-mixing of sorted nuclei or reads*

The *Loxodes magnus* MAC library had slightly more unique k-mers (2.0%) than the MIC library (1.3%), which may represent low-abundance contaminants, e.g. bacteria or feed algae.

To account for inadvertent cross-mixing during pooling of sorted nuclei or reads, two distinct sorted samples of each type of nucleus (and corresponding read libraries) were prepared per species. k-mer content was compared for each pairwise combination of libraries in each species. Pairwise comparison of individual replicates showed the same pattern as for the pooled libraries described above, both for *L. magnus* (Figure S3B) and *L. striatus* (Figure S4B), indicating that cross-mixing of nuclei types during pooling was unlikely.

#### *Genome assembly from short reads*

Illumina reads were trimmed to remove sequencing adapters, phage Phi X-174 sequence, and low-quality bases (Phred score < 28) with bbduk.sh from the BBTools software package v38.22 (http://sourceforge.net/projects/bbmap/), with options: ktrim=r qtrim=rl trimq=28 minlength=25. Trimmed reads were then assembled with SPAdes v3.13.0 [40] with options: --memory 380 -k 21,33,55,77,99,127.

#### *Genome assembly quality control*

Illumina genomic DNA read libraries were screened for SSU rRNA genes with phyloFlash v3.1,[41] using the SILVA 132 SSURef NR99 database,[42] to screen for potential contaminants. “Blob plots” of average contig coverage vs. contig GC% were plotted to visualize contig clusters that may represent contaminants. Most *Loxodes* contigs had ~25% GC, like most ciliate nuclear genomes; a contig cluster with ~45% GC was found to largely comprise low-complexity tandem repeats (Figure S6).

Completeness of the genome was estimated from proteins predicted in the *L. magnus* MAC genome by Pogigwasc (see main text Methods, “Gene prediction with Pogigwasc”), alongside 13 published ciliates and the alveolate *Perkinsus* marinus as outgroup (Table S4), with the Alveolata marker set in Busco v5.0.0,[43,44] using parameters: -m protein -l alveolata_odb10.

### *Simulation of k-mer comparisons for MIC and MAC of Paramecium tetraurelia*

*Paramecium tetraurelia* strain 51 MAC and MAC+IES genomes (v1.0) were obtained from ParameciumDB (https://paramecium.i2bc.paris-saclay.fr/files/Paramecium/tetraurelia/51/).[34] 10 million 150 bp paired-end reads were simulated from each library (random seed 0) with randomreads.sh from the BBTools software suite v38.18 (https://sourceforge.net/projects/bbmap/). A second MAC library was simulated with random seed 54321. Mixed libraries were simulated by subsampling the simulated libraries with reformat.sh from BBTools, using sample seed 12345 and sample rates 0.1, 0.3, 0.5, 0.7, and 0.9, then concatenating the respective subsamples in the required proportions. k-mer frequencies were compared with kat comp from Kat v2.4.2 [45] using k-mer length 19.

### *sRNA library preparation and sequencing*

Input RNA was quality assessed by capillary electrophoresis (small RNA chip, Agilent Bioanalyser); an Illumina-compatible NGS library was prepared with a NEXTflex Small RNA-Seq Kit v3 (Bioo Scientific) and sequenced in 2 x 150 bp paired-end read mode on an Illumina HiSeq3000 instrument at the Max Planck Genome Centre Cologne.

### *sRNA-seq analysis*

sRNA reads were trimmed with Trim Galore v0.6.7 (10.5281/zenodo.5127898) with parameters “--fastqc --small_rna --length 10 --paired”, merged with bbmerge [46] (parameters “mininsert=10 mininsert0=10 minoverlap=10), then mapped to the *Chlamydomonas reinhardtii* genome with bbmap (parameter: “k=10”) from BBTools v38.22.[47] Unmapped reads were mapped to the *Loxodes magnus* MAC and MIC genomes with bbmap and the same parameters. To generate length/5’ base composition histograms, replicates within the “fed” and “starved” conditions were combined since the individual replicate histograms were similar. Fed cell sRNAs appeared to be subject to less degradation of larger RNAs, as judged by the sRNA length histograms, and thus we focused on their properties.

Workflow: https://github.com/Swart-lab/loxodes-srna-workflow

#### *Transcriptome assembly*

For initial parametrization of the gene prediction model, a de novo transcriptome was assembled by mapping-guided assembly with Trinity v2.11.0,[48] using the preliminary SPAdes MAC genome assembly as a reference, with options: --SS_lib_type RF --genome_guided_max_intron 200.

Workflow: https://github.com/Swart-lab/loxodes-assembly-workflow

#### *RNA-seq count estimation*

GFF3 output from Pogigwasc was converted to GTF by GffRead,[49] then all “CDS” annotations were substituted with “exon” annotations. TPMCalculator [50] version 0.0.4 with optional switches “-c 14 -p -a” was then used to estimate TPM values across all the genes in the *Loxodes* MAC genome, using a BAM file of all the pooled RNA-seq from both starved and fed cells mapped to this genome.

#### *HiFi CCS read mapping to representative rDNA locus*

PacBio SMRT-seq HiFi CCS reads for the flow sorted MAC and MIC DNA were aligned to a complete representative rDNA sequence extracted from the *Loxodes magnus* MAC genome (as defined by a search of RFAM [51] using Infernal [52]) using pbmm2,[53] from the SMRT Link v12.0.0 software bundle from Pacific Biosciences (command ““pbmm2 align --preset CCS”; after indexing the reads with “pbindex”).

*Choice of 6mA detection threshold*

At the subread coverage threshold of 25 Pacific Bioscience's base modification detection software reported 1.13% of adenosines in the *L. magnus* MAC genome assembly as 6mA (5870562 potential 6mA positions out of 520983087 adenosines), compared to 0.0157% of adenosines in the MIC genome assembly as 6mA (98290 potential 6mA positions out of 626673708 adenosines). However, the modification identification quality values ($-10 \times \log_{10}$(probability of incorrect detection)) reported for the MIC genome in general were much lower than for the MAC genome (median 8 vs median 64). Thus, the percentage of 6mA bases called in the genomes was estimated at ≥ 25× coverage and an identification quality value ≥ 30.

Of the 32407 adenosines in a ~76 kb *L. magnus* mitochondrial contig (CAMPDZ020002550) co-assembled from residual DNA present in the MIC fraction, just one adenosine base was classified as 6mA at this threshold. This base was not followed by the characteristic thymine base and was called with an identification quality value of 5, and is likely thus a false positive modification call.
